# Permeation Across the Mycomembrane in Live Mycobacteria

**DOI:** 10.1101/2022.10.12.509737

**Authors:** Zichen Liu, Irene Lepori, Brianna E. Dalesandro, Mahendra D. Chordia, Taijie Guo, Karl Barry Sharpless, Jiajia Dong, M. Sloan Siegrist, Marcos M. Pires

## Abstract

The general lack of permeability of small molecules observed for *Mycobacterium tuberculosis* (*Mtb*) is most commonly ascribed to its unique cell envelope. More specifically, the outer mycomembrane is hypothesized to be the principal determinant for access of antibiotics to their molecular targets. Nonetheless, there is limited information on the types of molecular scaffolds that can readily permeate past the mycomembrane of mycobacteria. To address this, we describe a novel assay that combines metabolic tagging of the peptidoglycan scaffold, which sits directly beneath the mycomembrane, and a fluorescent labeling chase step, to measure the permeation of small molecules. We showed that the assay workflow was robust and compatible with high-throughput analysis in *Mycobacterium smegmatis* and *Mtb*. A small panel of molecules was tested and we found a large range in the permeability profile of molecules. Interestingly, the general trend is similar across the two types of mycobacteria, with some notable exceptions. We anticipate that this assay platform will lay the foundation for medicinal chemistry efforts to understand and improve uptake of both existing drugs and newly-discovered compounds into mycobacteria. The methods described, which do not require genetic manipulation, can be generally adopted to other species for which envelope permeability is treatment-limiting.

## Introduction

The tuberculosis (TB) pandemic continues to impact large swarths of the global population with an estimated one-third of the world population being latently infected with *Mycobacterium tuberculosis* (*Mtb*), the causative agent of TB. The health burden caused by TB is immense. Yearly, 1.5 million people die from TB infections and only approximately ∼50% of patients are successfully treated from multi-drug resistant TB.^1,2^ TB infections are inherently difficult-to-treat due to the low number of antimycobacterial agents that effectively clear the pathogen from infected patients. Similar to Gram-negative bacteria, mycobacteria possess an outer membrane (OM) that encases the entire cell (**Fig. 1A**).^3,4^ This membrane (mycomembrane) serves as a formidable barrier that is thought to hinder the penetration of small molecules (including antibiotics). As such, it has been implicated in endowing mycobacteria with a high level of intrinsic drug resistance to antimycobacterial agents.^5^

**Figure 1.**
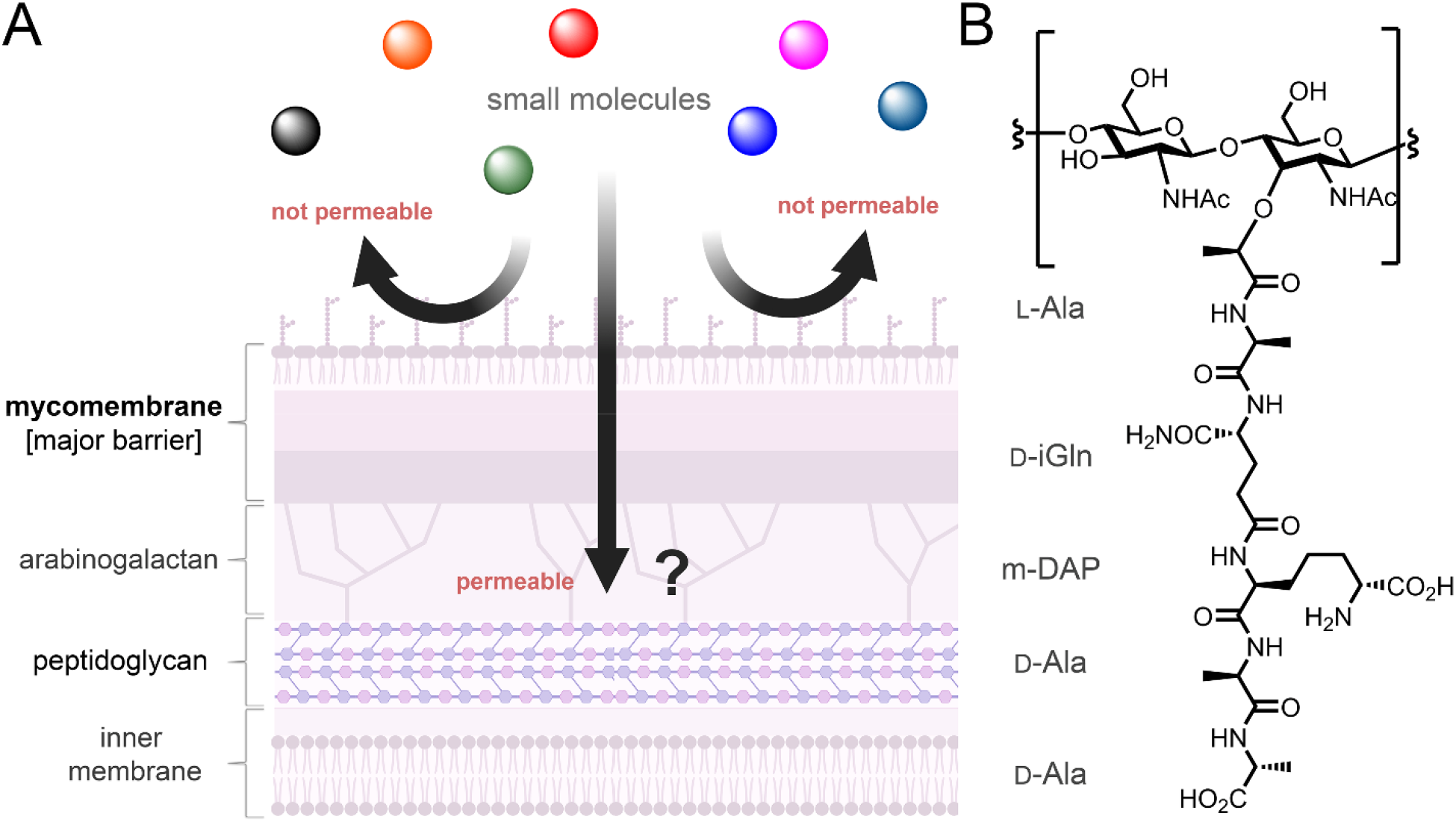
(**A**) Schematic representation of the cell surface of mycobacteria. The mycomembrane is generally purported to be the major barrier to the permeation of small molecules into mycobacteria. The peptidoglycan layer is localized beneath the mycomembrane, thus molecules that reach the peptidoglycan must have permeated through or across the mycomembrane layer. (**B**) Chemical structure of a canonical pentameric muropeptide of mycobacteria.

Given its location – lying at the interface between the potentially vulnerable inner components and the host – the mycomembrane is central to the host-mycobacteria relationship. Lack of permeation across the mycomembrane has long been hypothesized to be one of the primary reasons for the failure of antibiotics to accumulate in mycobacteria. The outer leaflet is composed primarily of trehalose dimycolate (TDM), which is conserved in all mycobacterial species, and intercalated by a number of noncovalently associated glycolipids, including some that are found only in pathogenic mycobacteria. The inner leaflet of the mycomembrane is comprised of arabinogalactan-esterified mycolic acids, the alkyl chains of which can range from C_60_ to C_90_.^6^ This waxy layer must pack tightly to fold the hydrocarbon chains into a prototypical 7-8 nm thick membrane.^7^ In doing so, the fluidity of the membrane is decreased^5^, thus also creating a less permeable barrier to most small molecules.

The therapeutic effectiveness of antimycobacterial agents depends on their ability to cross the mycomembrane to ultimately reach their cellular target. To this end, most antimycobacterial agents are relatively small and hydrophobic; antimycobacterial drugs that are larger have been proposed to cross *via* porins embedded within the mycomembrane. A prominent example of such a porin is MspA found in *Mycobacterium smegmatis* (*Msm*).^8^ In *M. tuberculosis*, however, there are no known Gram-negative-like porins in the mycomembrane, even though, an analogous role as transporter can be played by PE/PPE proteins^9-11^. Given the architecture of the mycobacterial cell wall, molecules that reach the peptidoglycan (PG) layer must have necessarily crossed the mycomembrane. Consequently, we reasoned that we could quantitatively probe the permeation of small molecules across the mycomembrane by quantifying the level of molecules that reach the PG scaffold.

PG is a mesh-like polymer made up of repeating disaccharides *N*-acetylglucosamine (GlcNAc) and *N*-acetylmuramic acid (MurNAc). Each MurNAc unit is connected to a short and unusual peptide (stem peptide) with the canonical sequence of L-Ala-D-iGln-*m*-DAP-D-Ala-D-Ala, in the case of *Mtb* (**Fig. 1B**).^12^ In all known bacteria, neighboring stem peptides are crosslinked by transpeptidases to endow the PG matrix with high levels of rigidity and integrity. Bacterial PG is an essential component of the bacterial cell wall, which makes its biosynthesis vulnerable to inhibition by small molecules such as β - lactams.^13,14^ We envisioned that site selective metabolic installation of a biorthogonal (“click chemistry”) handle within the PG scaffold of mycobacteria could be leveraged to assess the accumulation of small molecules across the outer mycomembrane. This workflow, <u>P</u>eptidoglycan <u>A</u>ccessibility <u>C</u>lick-<u>M</u>ediated <u>A</u>ssessme<u>N</u>t (PAC-MAN), could provide a platform to measure the permeation of any molecule that is modified with a small click handle. While a range of click chemistry handles have been developed to date that could potentially operate within the general workflow described, we focused on the combination of DiBenzoCycloOctyne (DBCO) and azide.^15,16^ This pair of reactive functional groups is biorthogonal and readily forms a stable triazole bond in the absence of metal catalysts based on strain promoted alkyne-azide cycloaddition (SPAAC).^17^ We expected that small molecules that readily permeate across the mycomembrane will react with PG-embedded DBCO; therefore, there will be fewer available DBCO epitopes to react with an azide-tagged fluorophore, thus resulting in a decrease in cellular fluorescence.

## Results and Discussion

We recently described a conceptually analogous assay that reports on the accessibility of larger biopolymers to and within the PG scaffold in *Staphylococcus aureus* (*S. aureus*).^18^ *S. aureus* were treated with an unnatural amino acid (D-lysine) modified with an ε -azide to label the entire PG scaffold with azide handles. Cells were then treated with fluorescently labeled biopolymers dually tagged – with DBCO and a fluorophore – to probe the accessibility of molecules to the cell surface. In the case of Gram-positive bacteria, the PG scaffold is fully exposed to the extracellular media and the biopolymers did not have to cross a membrane to be covalently anchored. Nonetheless, there are fundamental aspects of the assay that were adopted to establish PAC-MAN. In the case of PAC-MAN, we reasoned that the reactive handles needed to be reversed. The small molecules each included a compact azide tag that minimally perturbs the physiochemical properties of the test small molecules. In turn, the DBCO handle was conjugated to the PG *via* metabolic labeling that will display this reactive tag on the PG scaffold (**Fig. 2A**).

**Figure 2.**
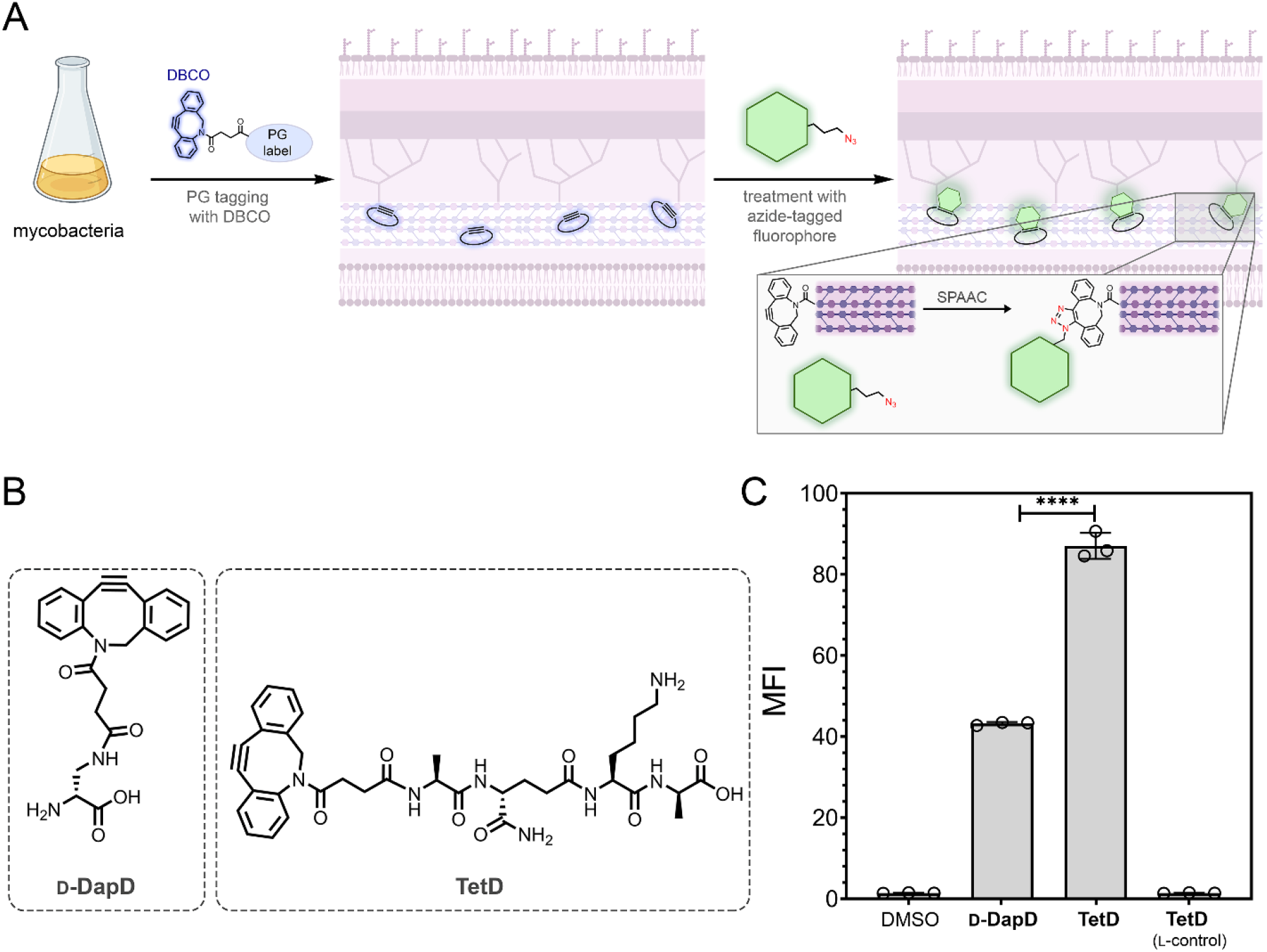
(**A**) Workflow schematic of the PG labeling with a DBCO-displaying agent followed by treatment with an azide-tagged fluorophore. (**B**) Chemical structures of **D-DapD** and **TetD**. (**C**) Flow cytometry analysis of *Msm* treated overnight with 25 μ M of synthetic PG analogs and incubated for 1 h with 25 μ M of **Fl-az**. Data are represented as mean +/- SD (n = 3). *P*-values were determined by a two-tailed *t*-test (* denotes a *p*-value□<□0.05, **□<□0.01, ***<0.001, ns□=□not significant).

To design the metabolic labels of mycobacterial PG, we considered that the DBCO unit could be linked to the side chain of a single D-amino acid. During cell growth, exogenous single D-amino acids supplemented in the culture medium are swapped in the place of the D-alanine that occupies the 4^th^ or 5^th^ position within the stem peptide in the PG layer.^3,19-34^ As an alternative, the DBCO could be linked to a synthetic mimic of the stem peptide. We^*28, 31*^, and others^*32-35*^, recently showed that synthetic analogs of PG stem peptides can be crosslinked into the growing PG scaffold of live cells, including that of *Mycobacterium smegmatis* (*Msm*) and *Mtb*. Generally, we had previously found that the *N*-terminus of the tetrapeptide is much more tolerant to large conjugates than the sidechain of single D-amino acids.^24^ For both types of PG metabolic tags, the inclusion of synthetic tags in the culture media during cellular growth results in the covalent incorporation of DBCO throughout the entire PG scaffold. Importantly, this tagging methodology is expected to be readily compatible with any mycobacteria (including drug-resistant strains and clinical isolates); therefore, PAC-MAN can be deployed against a number of mycobacterial species without the need for genetic manipulation.

We synthesized a single amino acid with a DBCO conjugated on the sidechain of D-Dap (**D-DapD**) and a tetrapeptide (**TetD**) with a DBCO conjugated on the *N*-terminus to empirically test the tolerance of the click handle on the metabolic label of live cells (**Fig. 2**). *Msm* cells were incubated with **TetD, D-DapD**, or vehicle to promote the tagging of the PG, washed to remove unincorporated tags, treated with azide-modified fluorescein (**Fl-az**), and cellular fluorescence was measured by flow cytometry (**Fig. 2A**). Satisfyingly, the background labeling of cells treated with vehicle was low, while cells treated with PG metabolic labels displayed considerably higher cellular fluorescence (**Fig. 2C**). As anticipated, the overall PG labeling levels observed with the single amino acid (**D-DapD**) were ∼40-fold above vehicle treated cells whereas labeling with **TetD** led to ∼85-fold fluorescence increase over vehicle. A diastereomeric version of **TetD** was synthesized in which the *C*-terminal D-Ala was switched to L-Ala. We had previously shown that transpeptidases are sensitive to the stereocenter on the *C*-terminus, which is the amino acid that is removed during the first step of transpeptidation.^29,35-37^ Similarly, fluorescence levels for *Msm* treated with the stereocontrol agent were similar to vehicle treated cells (**Fig. 2C**). These results establish that *Msm* labeling with **TetD** led to a higher signal-to-noise ratio, and, therefore, **TetD** became the primary PG tagging method.

The necessity of DBCO for the cellular labeling was tested by incubating *Msm* cells with fluorescein alone (lacking a biorthogonal reactive handle). Cellular fluorescence signals were found to be background levels in cells metabolically labeled with **TetD** followed with fluorescein (**Fig. 3A**). When the fluorescein was directly connected to the tetrapeptide (**TetFl**), fluorescence levels observed were nearly identical to cells labeled with **TetD** followed by azide-modified fluorescein (**Fl-az**), similar to our prior results in *Msm*.^29^ These results indicate that fluorescence levels observed with cells treated with the **TetD** followed by the azide-linked fluorophore are reflective of strain promoted alkyne-azide cycloaddition (SPAAC) reactions within the PG scaffold and that the SPAAC reaction was nearly complete.^15,17^ In agreement with the mechanism of incorporation, confocal microscopy of *Msm* treated with **TetD** followed by **Fl-az** revealed a labeling pattern that was consistent with cell surface labeling (**Fig. S1**).

**Figure 3.**
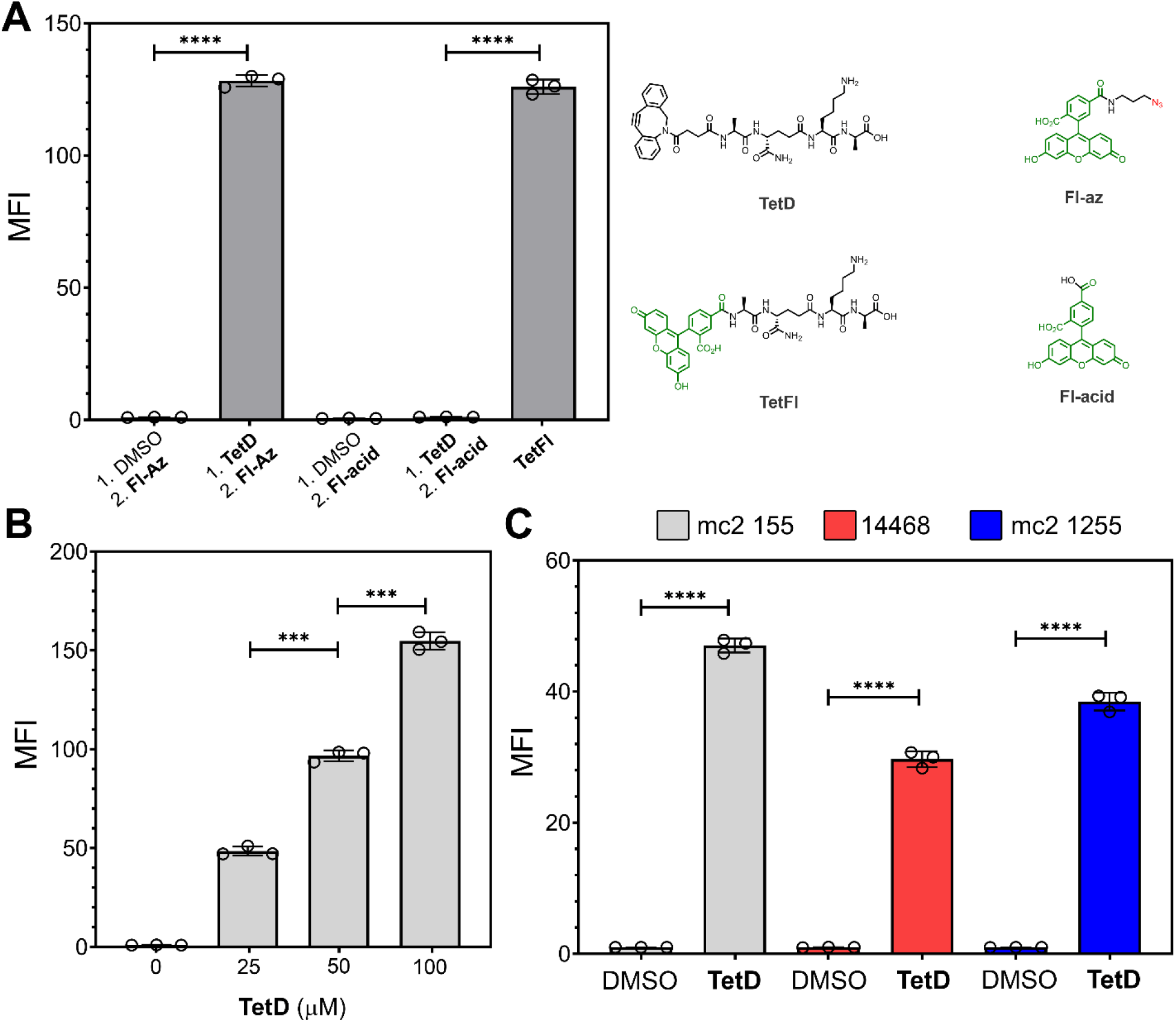
(**A**) *Left*: Flow cytometry analysis of *Msm* treated with 25 μ M of synthetic PG analogs and incubated with 50 μ M of **Fl-az**. Data are represented as mean +/- SD (n = 3). *Right*: Chemical structures of **TetD, TetFl, Fl-az**, and **Fl-acid**. (**B**) Flow cytometry analysis of concentration scan of **TetD** in *Msn* and incubated with 25 μ M of **Fl-az** for 1h at 37 °C. Data are represented as mean +/- SD (n = 3). (**C**) Different *Msm* strains (mc^2^ 155, ATCC 14468, mc^2^ 1255) were incubated with 25 μ M **TetD** then labeled with 50 μ M **Fl-Az** for 1h at 37 °C. Samples were then analyzed with flowcytometry. Data are represented as mean +/- SD (n = 3). *P*-values were determined by a two-tailed *t*-test (* denotes a *p*-value□<□0.05, **□<□0.01, ***<0.001, ns□=□not significant).

A number of parameters were tested next to optimize labeling conditions. A titration experiment was performed with **TetD** and a concentration-dependent increase in cellular fluorescence was observed from 25 to 100 μ M of **TetD** (**Fig. 3B**). Additionally, a concentration-dependent and time-dependent increase in cellular fluorescence was observed with varying concentration of **Fl-az** (**Fig. S2**). These results indicated that 50 μ M of **Fl-az** and an incubation period of 60 minutes led to a large increase in cellular fluorescence that should be sufficient for PAC-MAN. Understanding that the permeation of the fluorophore itself could be subject to the permeation barriers associated with the mycomembrane, we tested two additional azide-modified fluorophores: rhodamine 110 and coumarin (**Fig. S3**). While treatment of cells pre-labeled with **TetD** led to significant increases in cellular fluorescence with both fluorophores, they had lower signal-to-noise ratios relative to **Fl-az**. In the case of rhodamine 110, fluorescence levels were higher than **Fl-az**, albeit with a lower fold increase over unlabeled cells. These results likely reflect non-specific binding of rhodamine 110 to *Msm*. For coumarin, there was a large fold increase over untreated cells despite the lower brightness of the fluorophore, which may reflect its smaller size and its turn-on configuration. Two other *Msm* strains were found to be similarly labeled with a combination of **TetD** and **Fl-az**, an indication that the labeling strategy should be widely applicable to *Msm* (**Fig. 3C**). Combined, these results establish that the fundamental workflow of PAC-MAN (metabolic labeling followed by SPAAC with a fluorophore) can operate in the context of *Msm*.

A number of subsequent experiments were performed to establish the localization of the DBCO epitopes within the cell wall of *Msm*. Two of these experiments performed were specifically designed to test whether labeling levels were linked to PG transpeptidase activity, which are expected to crosslink the synthetic stem peptide mimic into the PG scaffold. A large number of β -lactam (e.g., penicillins, cephalosporins, and carbapenems) covalently inhibit PG transpeptidases, which should block the metabolic tagging by **TetD**.^38,39^ *Msm* cells were treated with increasing levels of meropenem in the presence of **TetD** and the fluorescence levels were analyzed as before. Meropenem is known to inhibit L,D-transpeptidases (Ldts) that are expected to be the mediators of **TetD** incorporation. A concentration dependent decrease in cellular fluorescence was observed, which is consistent with transpeptidase processing of **TetD** (**Fig. 4A**). We next sought a genetic approach to delineate the labeling mechanism. An Ldt-deletion *Msm* strain, which has 5 of the Ldts genetically deleted (*ldt*Δ*5*)^29,40^, was likewise treated with **TetD**. Relative to the parental cells, fluorescence levels decreased ∼7-fold and are consistent with metabolic processing by Ldts (**Fig. 4B**). Fluorescence levels were found to be higher than cells treated with meropenem, which may indicate that there remains **TetD** processing in *ldt*Δ*5*. Given that the third position on the synthetic peptide contains a lysine group, it is possible that some of the PG incorporation is also mediated by Penicillin Binding Proteins (PBPs) as acyl-acceptor strands.

**Figure 4.**
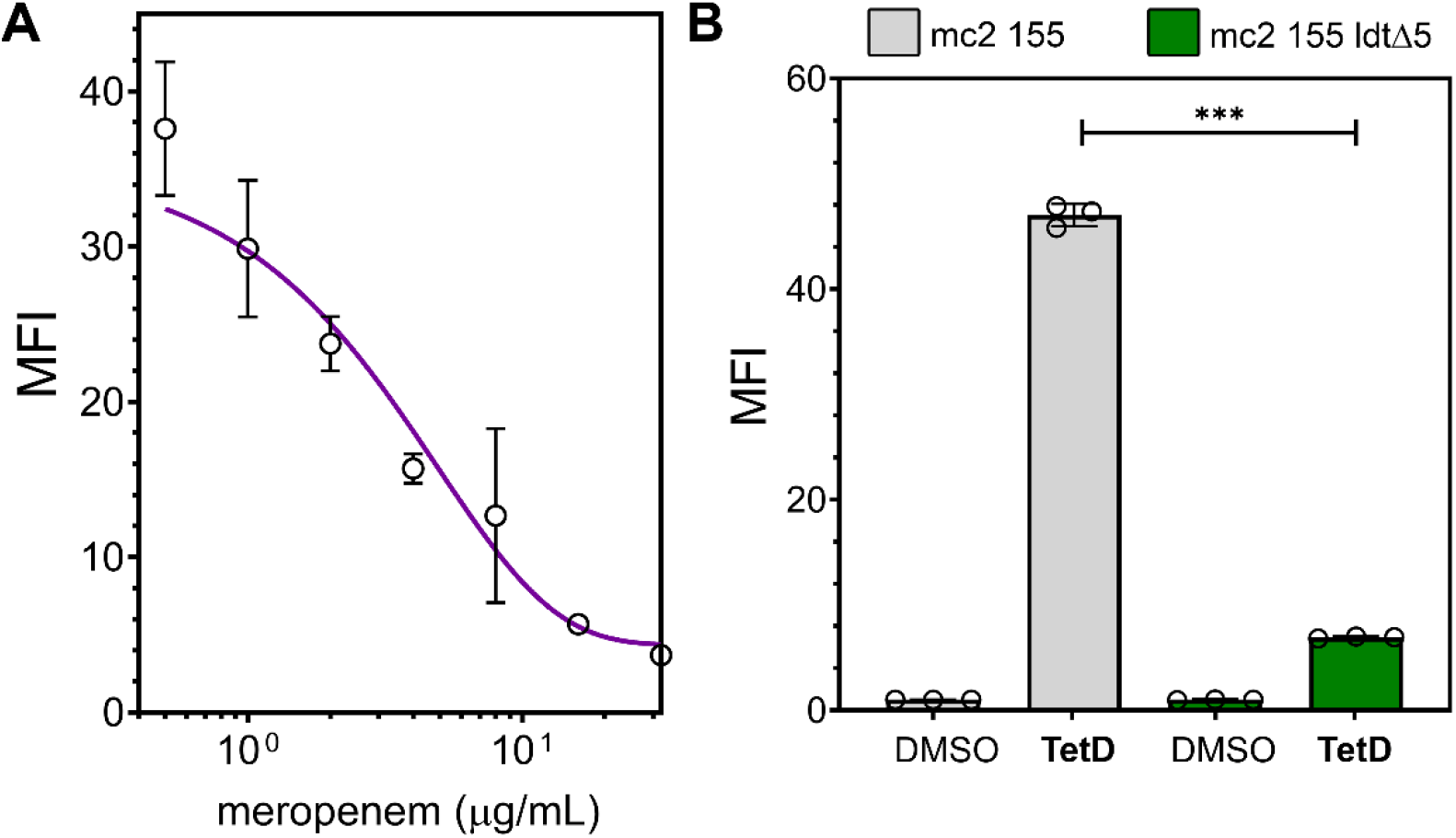
(**A**) Flow cytometry analysis of meropenem inhibition of incorporation of **TetD**. *Msn* was incubated with 25 μ M **TetD** and different concentration of meropenem then labeled with 50 μ M **Fl-az** for 1h at 37 °C. (**B**) *Msn* mc^2^ 155 and the Ldt-null strain, *ldt*Δ 5, were incubated with 25 μ M **TetD** then labeled with 50 μ M **Fl-az** for 1 h at 37 °C. Samples were then analyzed with flow cytometry. Data are represented as mean +/- SD (n = 3). *P*-values were determined by a two-tailed *t*-test (* denotes a *p*-value□<□0.05, **□<□0.01, ***<0.001, ns□=□not significant).

Further confirmation of the localization of the PG tag was obtained using LC-MS analysis. Sacculi from *Msm* cells that had been labeled (**TetD** followed by **Fl-az**) were isolated using standard protocols^41^ and digested with mutanolysin to yield muropeptides. These PG fragments were then analyzed by mass spectrometry. Transpeptidation reactions would be expected to generate dimeric (or larger oligomers) muropeptides containing the synthetic stem peptide mimic. Our results confirmed the presence of dimeric species crosslinked in a 3-3 configuration, which is consistent with Ldt processing (**Fig. S4**). Confocal microscopy of whole cells and isolated sacculi were further used to delineate the localization of the fluorescent signal (**Fig. S5**). Additionally, we used a protocol we recently described (SaccuFlow)^35^ to establish localization of the fluorescent signal within the mycobacterial cell wall. Starting with the intact cell wall metabolically labeled and reacted with **Fl-az**, individual layers were progressively removed and we found that cells tagged with **TetD** retained the fluorescence signal through the final step, which should yield only the PG (**Fig. 5**). In contrast, *Msm* cells metabolically labeled with a trehalose^42-45^ based tag (2-TreAz), which becomes noncovalently embedded within the mycomembrane, results in a minimal fluorescence of the isolated PG layer. Overall, these results are consistent with the installation of DBCO exclusively within the PG of *Msm* and establish that DBCO can be used as a reporter of molecules crossing the mycomembrane.

**Figure 5.**
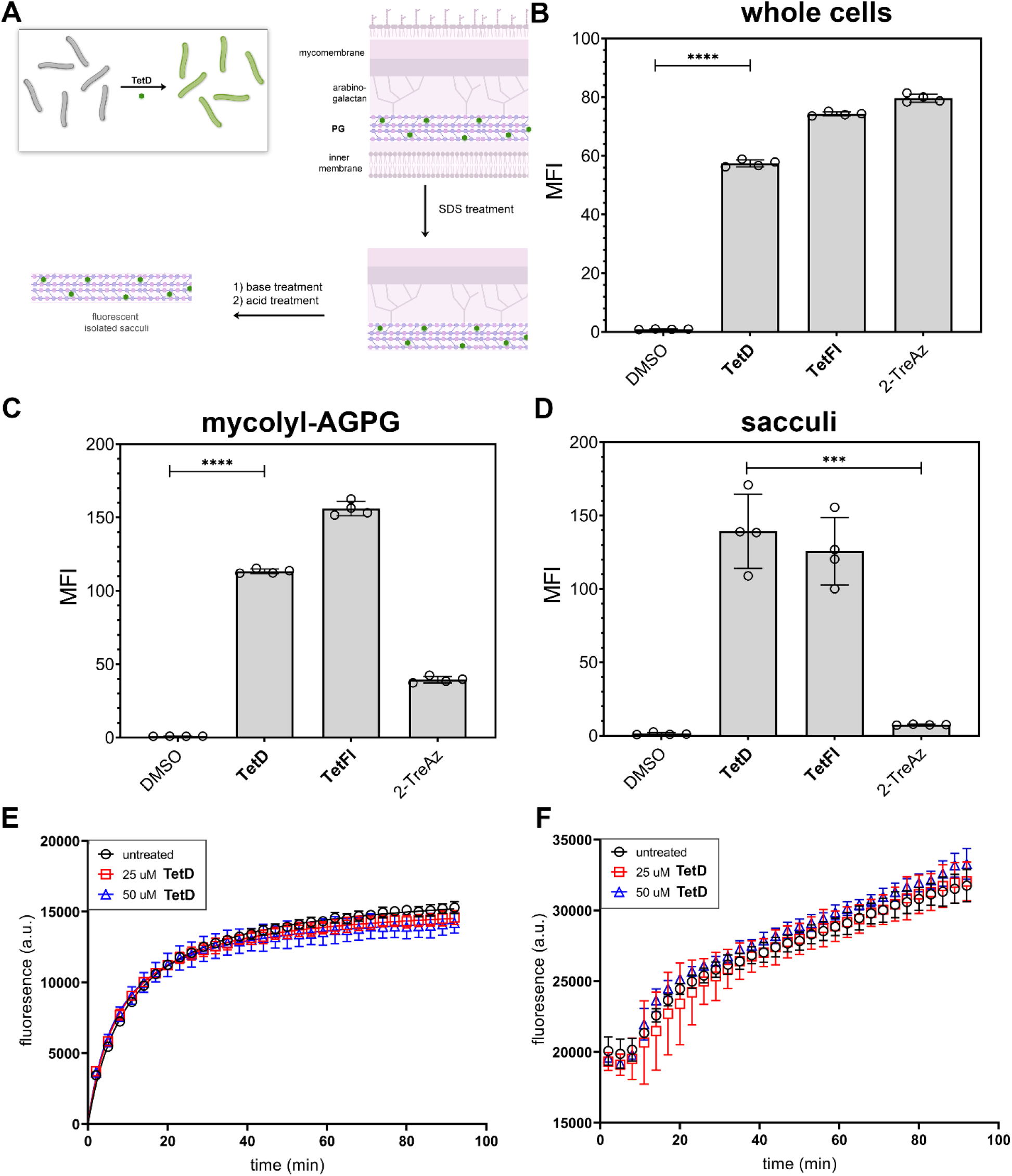
(**A**) Schematic representation of the steps in the workflow from labeled whole cells to isolated sacculi. Flow cytometry analysis of *Msm* mc^2^ 155 cells that were incubated with 25 μ M followed by treatment with 50 μ M of **Fl-az**. Analysis was performed with intact whole cells (**B**), mycolyl-AGPG (**C**), and isolated sacculi (**D**). Data are represented as mean +/- SD (n = 4). *P*-values were determined by a two-tailed *t*-test (* denotes a *p*-value□<□0.05, **□<□0.01, ***<0.001, ns□=□not significant). (**E**) Ethidium Bromide accumulation analysis. *Msm* mc^2^ 155 cells were incubated with 25 μ M or 50 μ M **TetD** before dispensing into a Costar 96-well black opaque flat bottom plate with 5 μ M ethidium bromide in triplicate. The fluorescence intensity was measured using a Synergy H1 microplate reader for 90 min with 3 min intervals (excitation 530 nm, emission 590 nm). (**F**) Nile red accumulation analysis. *Msm* mc^2^ 155 cells that were incubated with 25 μ M or 50 μ M **TetD** before dispensing into a Costar 96-well black opaque flat bottom plate with 10 μ M Nile red in triplicate. The fluorescence intensity was measured using a Synergy H1 microplate reader for 90 min with 3 min intervals (excitation 540 nm, emission 630 nm).

A series of experiments were performed to determine the impact of **TetD** on the cell wall of *Msm*. To start, the overall cellular viability of tagged cells was verified by treating *Msm* with increasing concentrations of **TetD** and cells were enumerated by colony forming unit (CFU) analysis and additionally analyzed by optical density measurements (**Fig. S6**). CFU count remained unchanged, an indication that there is no observable change to *Msm* viability. Similarly treated cells were also visualized by phase contrast and cellular structure appeared unaltered relative to vehicle-treated cells (**Fig. S7**). To directly probe the integrity of the mycomembrane, cells were analyzed for ethidium bromide (EtBr) accumulation/efflux and Nile red uptake by fluorescence measurements.^46-50^ These dyes have been previously used as indicators of mycomembrane integrity as disruption to the mycomembrane will result in increased intracellular accumulation of these agents, which leads to higher cellular fluorescence. *Msm* cells were treated with vehicle or **TetD**, washed, stained with EtBr (hydrophilic dye) or Nile Red (hydrophobic dye), and whole cell accumulation was measured using a fluorescence plate reader (**Fig. 5E** and **5F**). The results showed that in both cases, there was no significant change in the accumulation of the permeability probe when cells were co-incubated with **TetD**. Taken together, these results confirm that *Msm* cells whose PG were metabolically tagged by treatment with **TetD** retain their cellular viability and, importantly for downstream assays, the permeability profile of mycomembrane is not disrupted.

Next, we set out to benchmark the ability of the assay to report on the permeability of a set of test small molecules by performing a pilot screen (**Fig. 6A**) with azide-conjugated compounds not only in *Msm* but also in *Mtb*. These molecules cover a range of chemical space with a mixture of aliphatic and aromatic features and also a range of net charges (**Fig 6B**). To test whether the DBCO-azide reaction was complete within the timescale of the assay, we modified amine-terminated beads to display DBCO epitopes (**Fig. S8**). The beads are compatible with flow cytometer analysis, thus providing a similar workflow as the live cell analysis but without the mycomembrane permeability barrier. Our results showed that the molecules within the azide-modified panel readily react.

**Figure 6.**
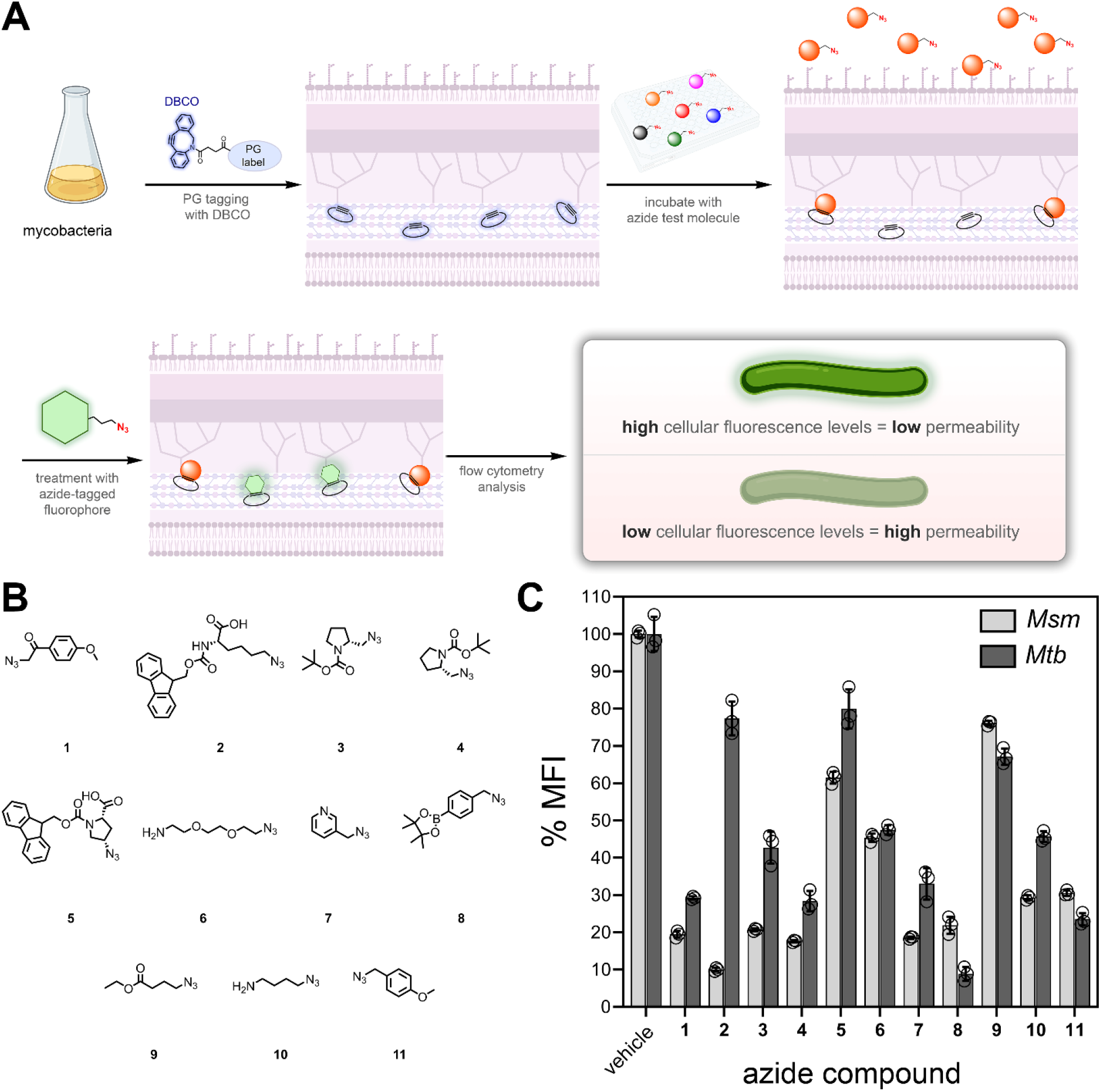
(**A**) Schematic representation of PAC-MAN. First, mycobacterial cells were metabolically tagged with **TetD**, followed by an incubation period with the test molecule modified an azide, cells were then incubated with **Fl-Az**, and finally flow cytometry analysis was performed. (**B**) Chemical structures of the initial panel of azide-modified molecules that were tested. (**C**) Flow cytometry analysis of *Msm* and *Mtb. Msm* were treated for 14 h (∼5 generations) with 25 μ M of **TetD**, incubated for 2 h with 50 μ M of each indicated test molecule, washed, and incubated with 50 μ M of **Fl-az** for 1 h at 37 °C. Signals were normalized to DMSO. (**D**) Flow cytometry analysis of PAC-MAN assay performed on Δ*leuD* Δ*panCD Mtb. Mtb* were incubated for 72 h with 25 μ M TetD (∼2 generations), washed, and incubated for 2 h with 50 μ M of each test molecule. Bacteria were then centrifuged to remove the test molecules and treated with 50 μ M of **FL-az** for 1 h. Signal was normalized using the bacteria not pre-incubated with **TetD** sample as 0% and the DMSO-treated bacteria as 100%. Data are represented as mean +/- SD (n = 3). *P*-values were determined by a two-tailed *t*-test (* denotes a *p*-value□<□0.05, **□<□0.01, ***<0.001, ns□=□not significant).

With the general workflow of PAC-MAN optimized, we set out to evaluate a panel of small molecules for their ability to accumulate past the mycomembrane. After confirming both the tolerability of **TetD** probe and its effectiveness of labeling in *Mtb* (**Fig. S9**), we proceeded with a pilot screen of 11 azido-compounds. The pilot screen was performed to demonstrate the robustness of the assay in a 96-well plate format and to characterize its reproducibility. PAC-MAN was initiated by metabolically tagging the PG of mycobacteria upon treatment with **TetD** (**Fig. 6A**). Batch treatment ensures similar PG labelling across the entire cell population. Cells were then washed to remove unincorporated molecules and bacterial cells and were dispensed into individual wells in a 96-well plate. Each well contained a unique azide-modified small molecule. For molecules that permeate past the mycomembrane, they were expected to covalently react with the PG scaffold *via* SPAAC. Next, cells were treated with **Fl-az** to quantify the level of unreacted DBCO epitopes.^51^ Finally, cells were analyzed by flow cytometry to measure total cellular fluorescence.

Screening 11 azide-modified test molecules revealed that even within this small panel of molecules there was a significant dynamic range in signal both for *Msm* and *Mtb* (**Fig. 6C**). The small aromatic molecule **1** displayed high apparent permeation in both types of mycobacteria. For compound **2**, there was considerable apparent permeation across the mycomembrane in *Msm* but this was not the case of *Mtb*. This could indicate that the hydrophobicity could have a variable impact in permeation across the mycomembrane. Unexpectedly, our results show one example of compound with higher apparent permeability through *Mtb* OM compared to *Msm* (compound **8**). Since *Mtb* mycomembrane is generally recognized as highly impermeable, especially compared to that of *Msm*, understanding the mechanism underlying this and similar phenomena, could possibly lead to the discover of transporters that are present in *Mtb* but not of *Msm*. A concentration dependent analysis of **2** and **6** in *Msm* revealed that it had an EC_50_ of apparent permeation of ∼6 μ M and ∼60 μ M, respectively (**Fig. S10**).

The initial panel of azides was informative in terms of the reproducibility and it provided the first direct comparison between *Msm* and *Mtb*. To further demonstrate the applicability of PAC-MAN, we subjected *Msm* to four azide-modified molecules that were drug-like in their chemical composition (**Fig. 7**). These molecules were generated from their commercially available primary amine precursor using the recently described diazotizing reagent.^52^ From these results, sulfonamide displayed the lowest EC_50_ of apparent permeation of ∼4.5 μ M while pazufloxacin had EC_50_ ∼22 μ M. The two other azide-modified molecules showed minimal apparent accumulation across the concentrations tested. Furthermore, 2 of the test conditions (a positive control with a readily permeable molecule and a negative control treated with PBS) were tested in *Msm* across an entire 96-well plate (**Fig. S11**) and it had a Z’ value of 0.89. There was a high level of stability observed, which indicates that PAC-MAN can be readily expanded to a high-throughput format. Critically, these results confirm that PAC-MAN can be used to compare the apparent permeation of molecules in mycobacteria.

**Figure 7.**
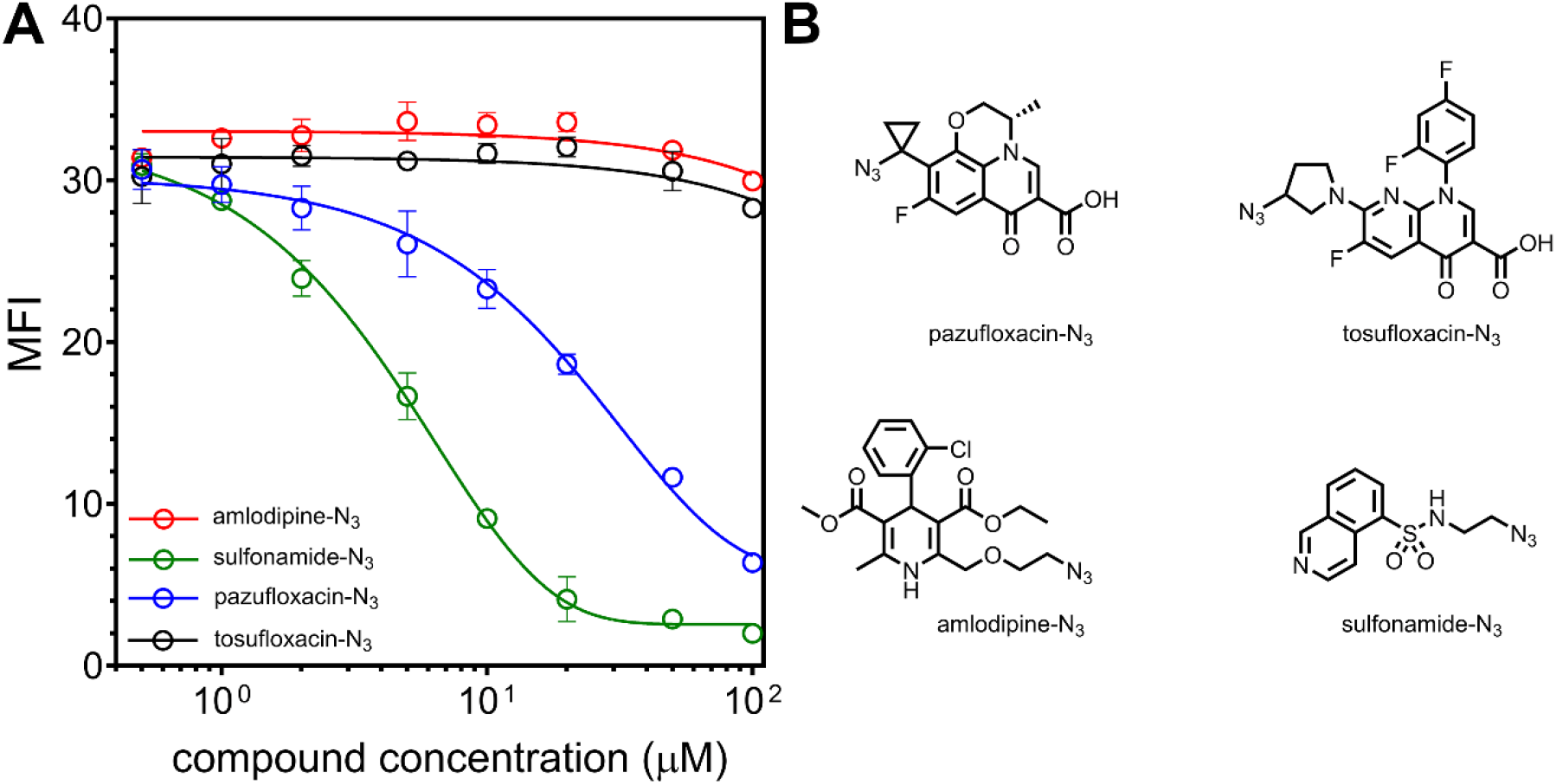
(**A**) Flow cytometry analysis of apparent permeation in *Msm. Msm* were treated for 14 h (∼5 generations) with 25 μ M of **TetD**, incubated for 2 h with varying concentrations of each indicated test molecule, washed, and incubated with 50 μ M of **Fl-az** for 1 h at 37 °C. Signals were normalized to DMSO. Data are represented as mean +/- SD (n = 3). (**B**) Chemical structures of four of azide-modified molecules that were tested.

## Conclusion

Permeability of small molecules into mycobacteria has for long been proposed to be primarily dictated by their (in)ability to cross the outer most membrane bilayer, the mycomembrane. The unique composition and physiochemical features of the mycomembrane can result in restricted passive permeation. Mycobacteria generally lack Gram-negative-like porins that facilitate the accumulation of polar molecules. This is especially the case for *Mtb* for which there are no known porins that have been show to promote the permeation of small molecule antibiotics, even if an analogous role can be played by PE/PPE proteins^9^. We have established an assay (PAC-MAN) that leverages the metabolic installation of biorthogonal epitopes within the PG layer to robustly determine the ability of molecules to permeate past the mycomembrane. Molecules of interest are tagged with a small azide tag and when they reach the PG layer they react with DBCO, thus imprinting their permeation. A chase step with an azide-linked fluorophore is performed to reveal the level of available DBCO epitopes. The assay was optimized to develop the baseline workflow conditions. Next, we showed with a panel of azide-tagged molecules that permeation profiles could be reproducibly performed in *Msm* and pathogenic *Mtb*. We anticipate that PAC-MAN could form the basis of an assay platform that could reveal permeation profiles of azide-tagged molecules in a parallel fashion. In this format, it should be possible to reveal biological contexts (*e*.*g*., inside macrophages, in more physiologically relevant culture conditions, in the presence of small molecule adjuvants) that promote or disfavor the permeation of molecules across the mycomembrane.

## Acknowledgement

This study was supported by the NIH grant GM124893-01 (M.M.P.), NIH DP2 AI138238 (M.S.S.), and the University of Massachusetts Amherst Institute for Applied Life Sciences Midigrant (I.L.). *ldt*Δ*5* was a kind gift from M. Pavelka (University of Rochester). We thank Dr. Amy Burnside, director of the University of Massachusetts Amherst Flow Cytometry Core facility, for technical support.

## Supporting Information

Additional figures, tables, and materials/methods are included in the supporting information file.

