## Supporting Information for "Permeation Across the Mycomembrane in Live Mycobacteria"

| **TABLE OF CONTENTS** | |
| --- | --- |
| **SUPPORTING FIGURES** | **S3-S13** |
| Figure S1. Confocal of *M. smegmatis* labeled with **TetD** and **Fl-Az** | S3 |
| Figure S2. Labeling kinetics with **Fl-Az** | S4 |
| Figure S3. Labeling of *M. smegmatis* with different dyes | S5 |
| Figure S4. Mass analysis of stem peptide from labeled *M. smegmatis* | S6 |
| Figure S5. Confocal of sacculi from labeled *M. smegmatis* | S7 |
| Figure S6. CFUs of labeled *M. smegmatis* | S8 |
| Figure S7. Phase contrast imagine of *M. smegmatis* labeled with **TetD** | S9 |
| Figure S8. Synthesis diagram and test molecules competition on DBCO coated polystyrene beads | S10 |
| Figure S9. Viability and labeling of *M. tuberculosis* | S11 |
| Figure S10. EC50 curve of selected test molecules **2** and **6** on *M. smegmatis* | S12 |
| Figure S11. High throughput test run in 96-well plate format | S13 |
| **MATERIALS AND METHODS** | **S14-S31** |
| General materials resources table | S14 |
| General Chemistry methods and Instruments | S15 |
| Synthesis and characterization of TetD | S15-S18 |
| Synthesis and characterization of TetD(L control) | S19-S21 |
| Synthesis and characterization of D-DapD | S21-S24 |
| Biological methods | S25-S30 |
| **REFERENCES** |  |

**Supporting Figures**

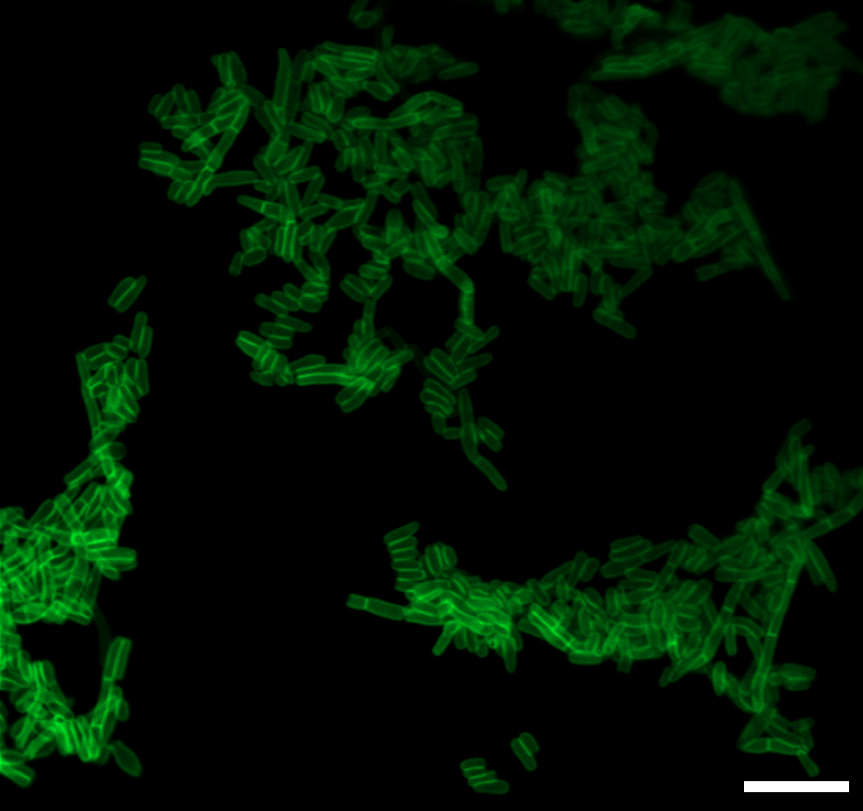

**Figure S1.** Confocal microscopy analysis of *M. smegmatis (Msn)* metabolically labeled with 100 μM **TetD** then labeled with 100 μM **Fl-Az** for 1 h at 37 ℃. Scale bar = 5 μm.

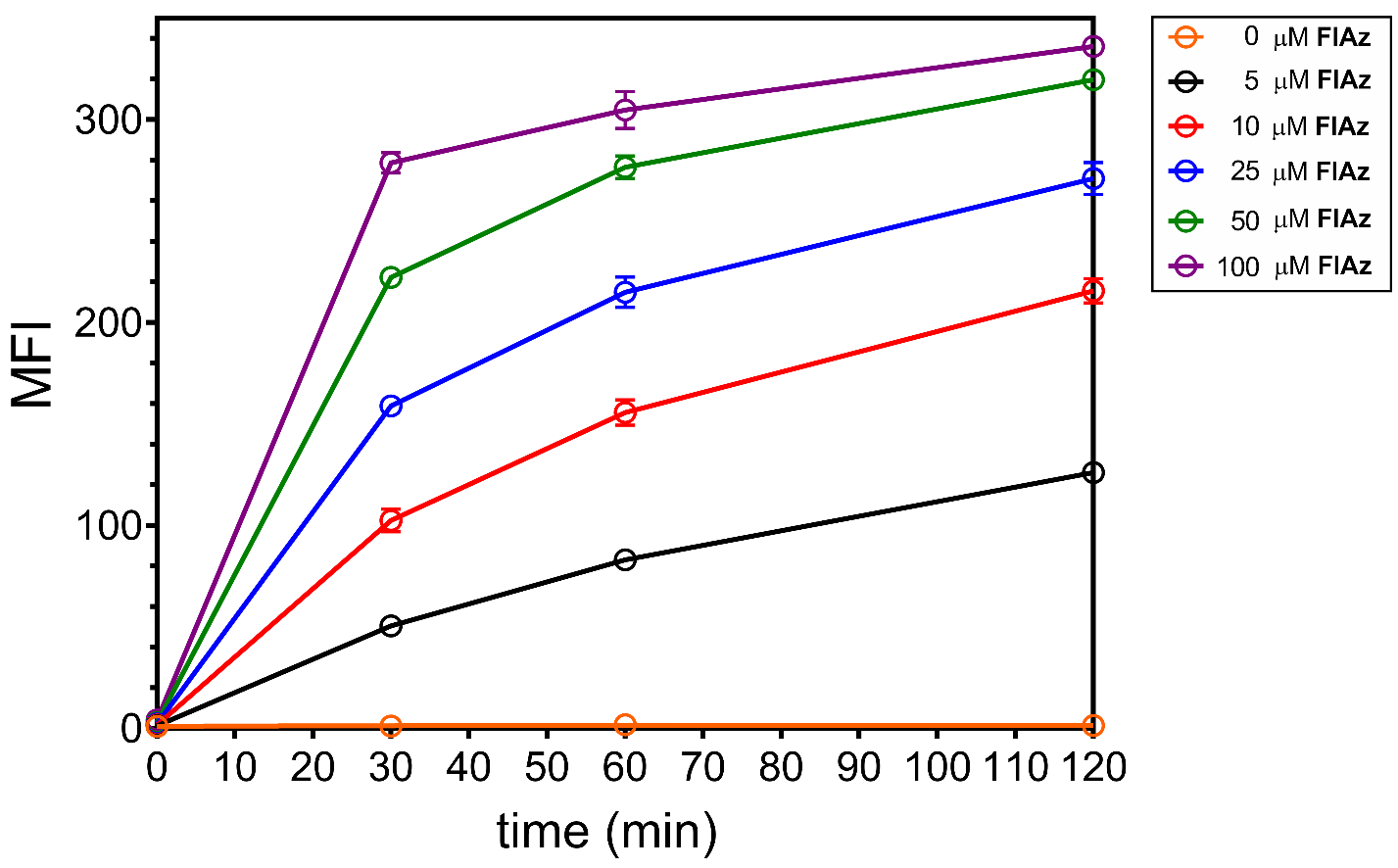

**Figure S2.** Flow cytometry analysis of labeling kinetics of different concentration of **Fl-az**. *Msm* was incubated with 25 μM **TetD** then labeled with **Fl-az** for different time lengths at 37 ℃. Samples were then analyzed with flowcytometry. Data are represented as mean +/- SD (n = 3).

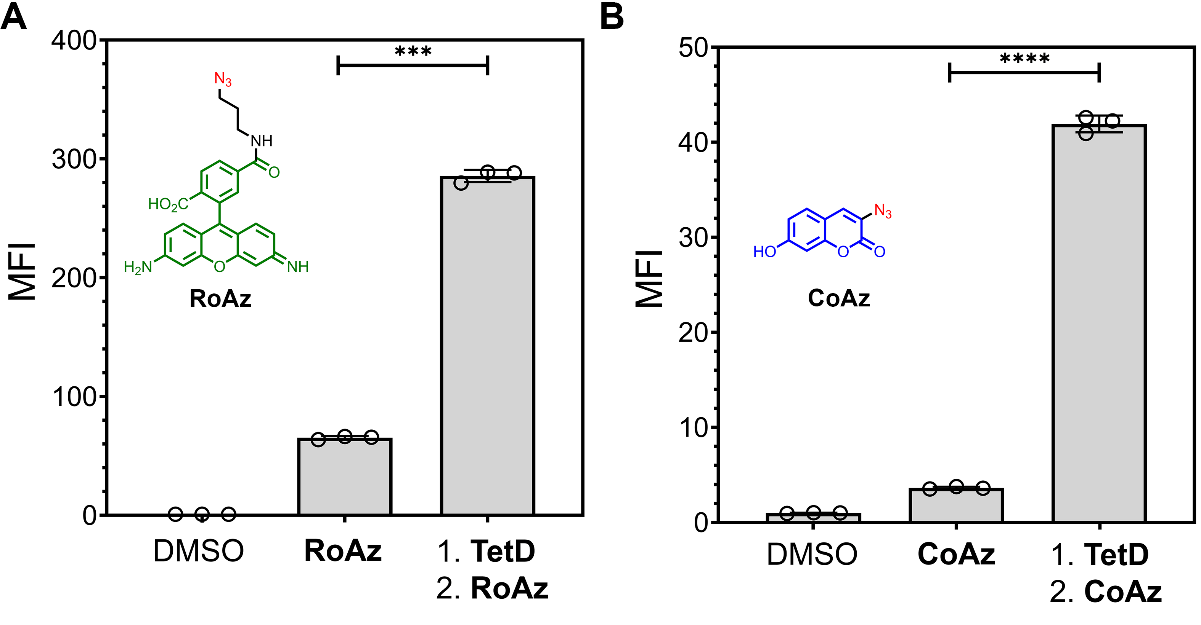

**Figure S3.** Flow cytometry analysis of *Msn* labeled with different dyes. **RoAz**, 6-azido- rhodamine 110. **CoAz**, 3-Azido-7-hydroxy coumarin. *Msn* mc^2^ 155 was incubated with 25 μM **TetD** then labeled with 50 μM **RoAz** or **CoAz** for 1h at 37 ℃. Data are represented as mean +/- SD (n = 3). *P*-values were determined by a two-tailed *t*-test (* denotes a *p*-value < 0.05, ** < 0.01, ***<0.001, ns = not significant).

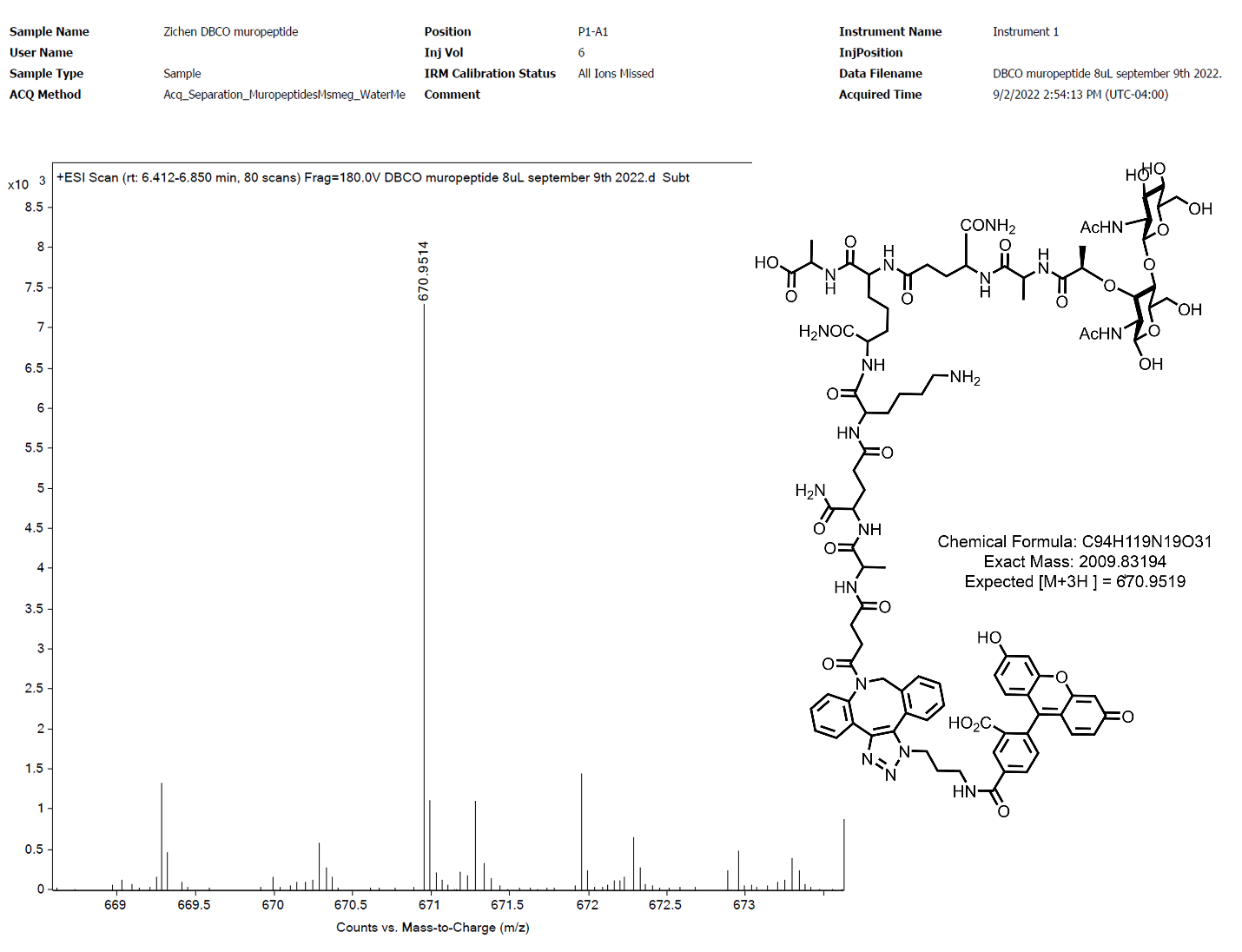

**Figure S4.** Mass spectrometry analysis of stem peptides from *Msn* incubated with 25 μM **TetD** and then labeled with 50 μM **Fl-Az** for 1h at 37 ℃. After incubation, the cells were harvested, with the peptidoglycan isolated and digested with mutanolysin and lysozyme, and the supernatant was analyzed by LC-MS.

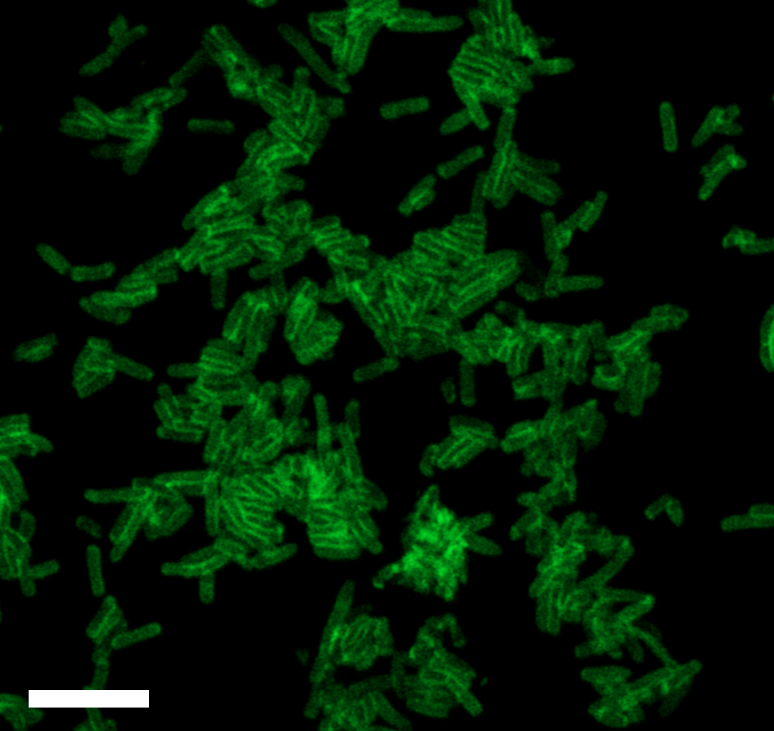

**Figure S5.** Confocal microscopy analysis of sacculi from *Msn* metabolically labeled with 25 μM **TetD** then labeled with 50 μM **Fl-Az** for 1 h at 37 ℃. Scale bar = 5 μm.

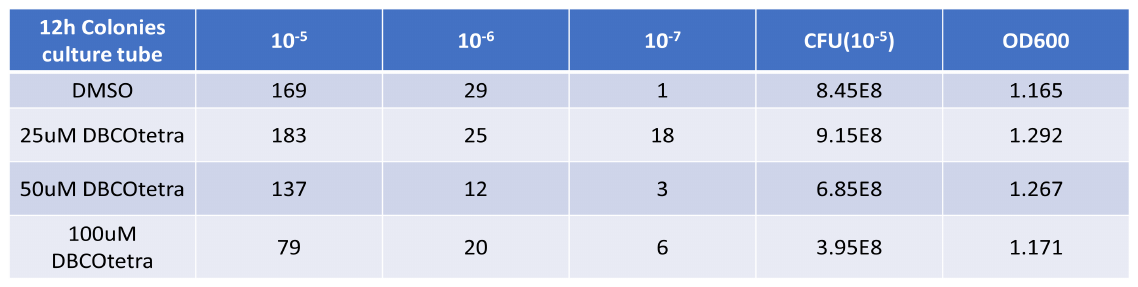

**Figure S6.** Colony forming unit analysis and OD_600_ measurements were performed in *Msn* mc^2^ 155 cells treated with varying concentrations of **TetD**.

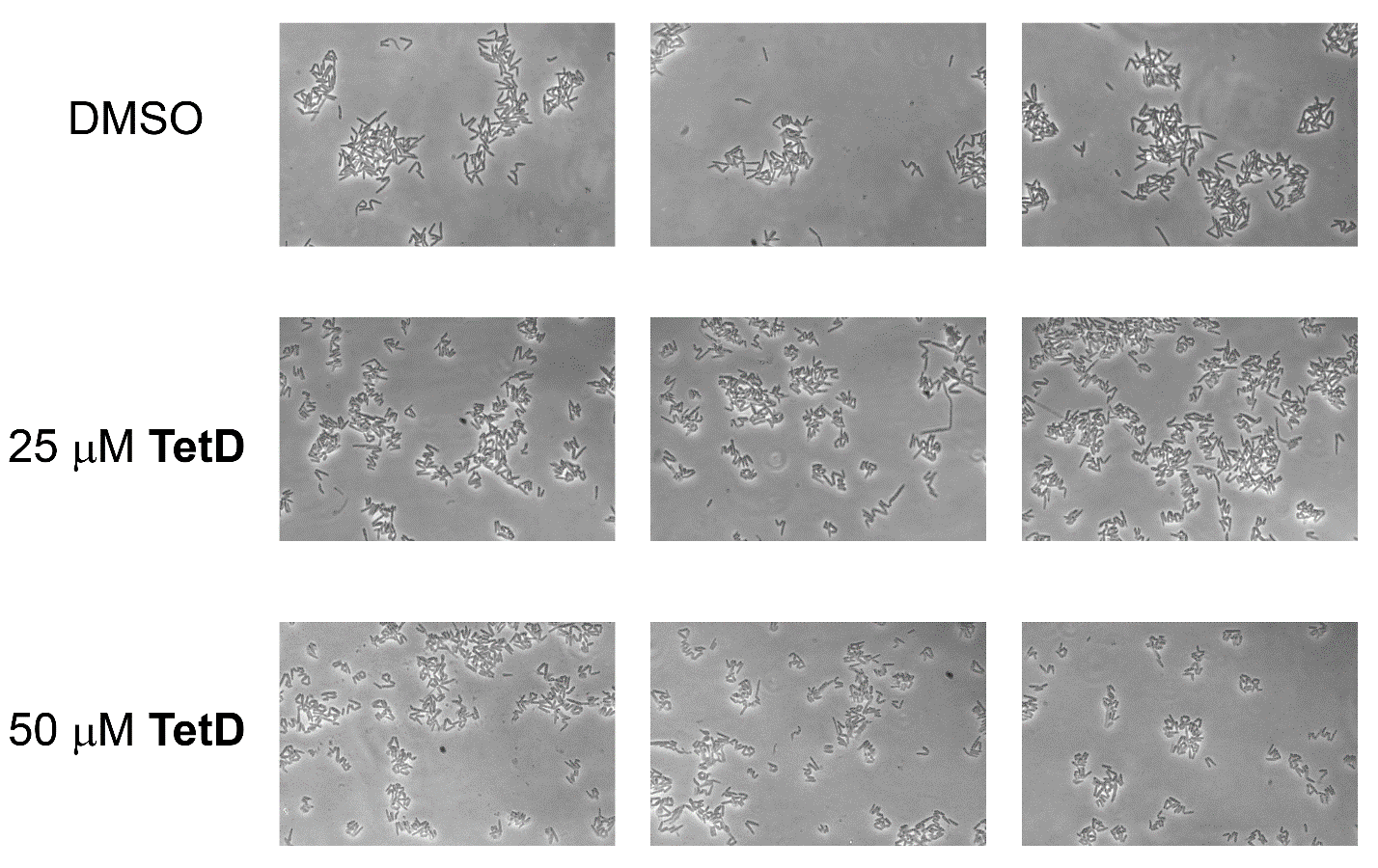

**Figure S7.** Phase contrast imaging of *Msn* mc^2^ 155 cells treated with varying concentrations of **TetD**.

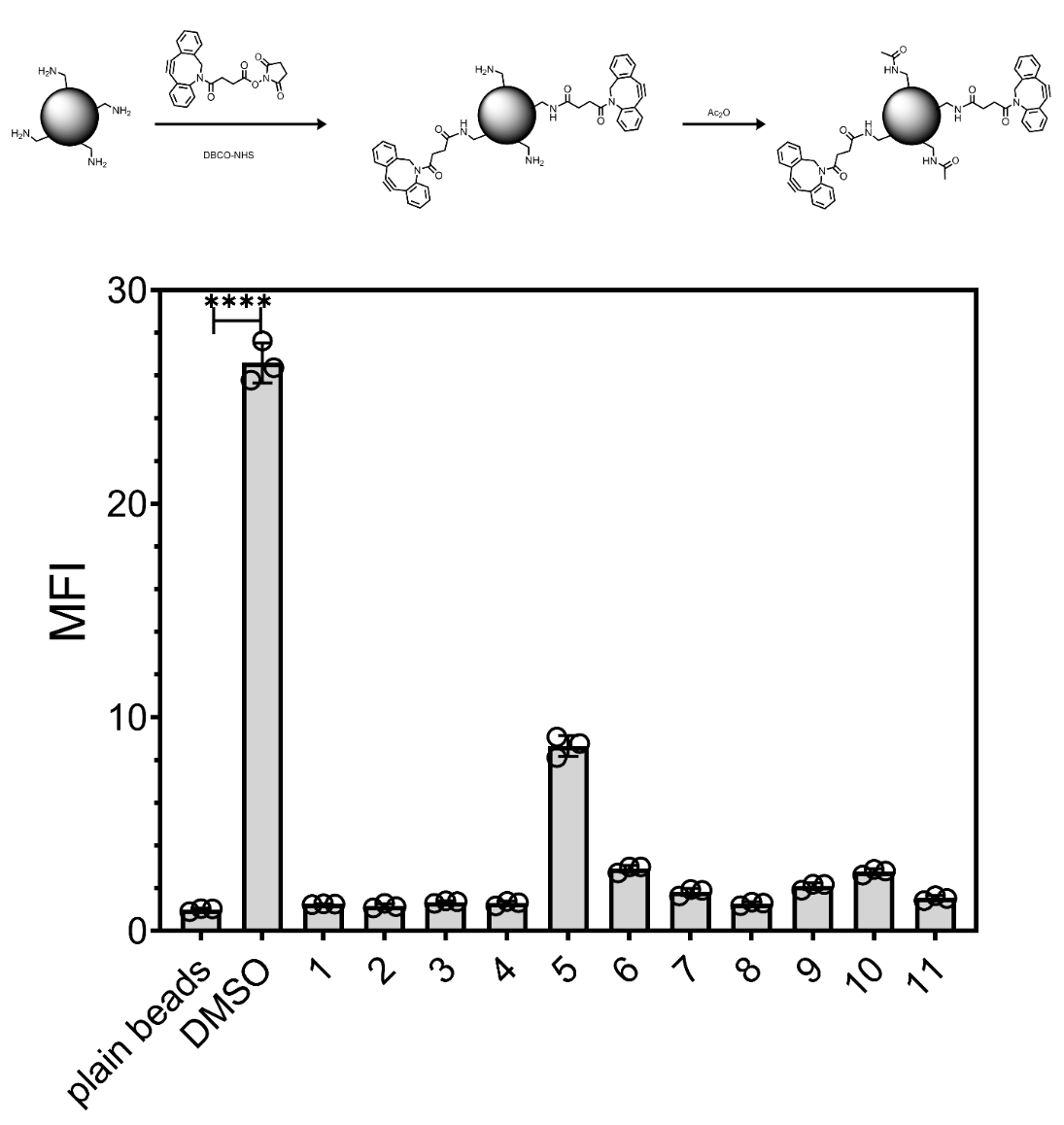

**Figure S8.** Amino terminated polystyrene beads were reacted with NHS-activated DBCO. Free amines that remained unreacted were capped with acetic anhydride. DBCO-modified beads were incubated with the panel of 50 μM azide-modified molecules 2h at 37 ℃ in PBS, followed by 1h incubation with 50 μM **Fl-az**, and analyzed by flow cytometry.

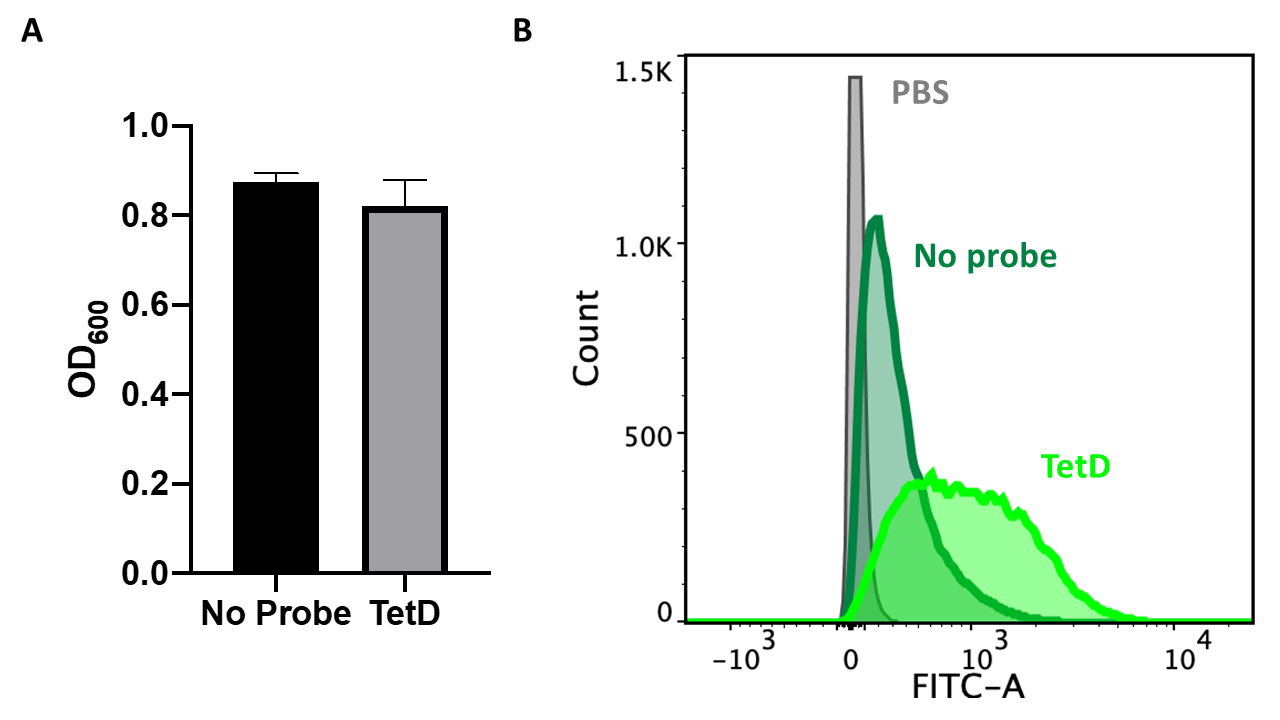

**Figure S9.** Δ*leuD* Δ*panCD* *M. tuberculosis* labeling using **TetD** peptidoglycan-targeting probe. (**A**) Bacteria viability after 36 h of incubation with 25 μM **TetD** measured by OD_600_. (**B**) Flow cytometry analysis of bacteria after 72 h of incubation with 25 μM **TetD** and subsequent labeling with **FL-az** 50 μM for 1h. Data are represented as mean +/- SD (n = 3). *P*-values were determined by a two-tailed *t*-test (* denotes a *p*-value < 0.05, ** < 0.01, ***<0.001, ns = not significant).

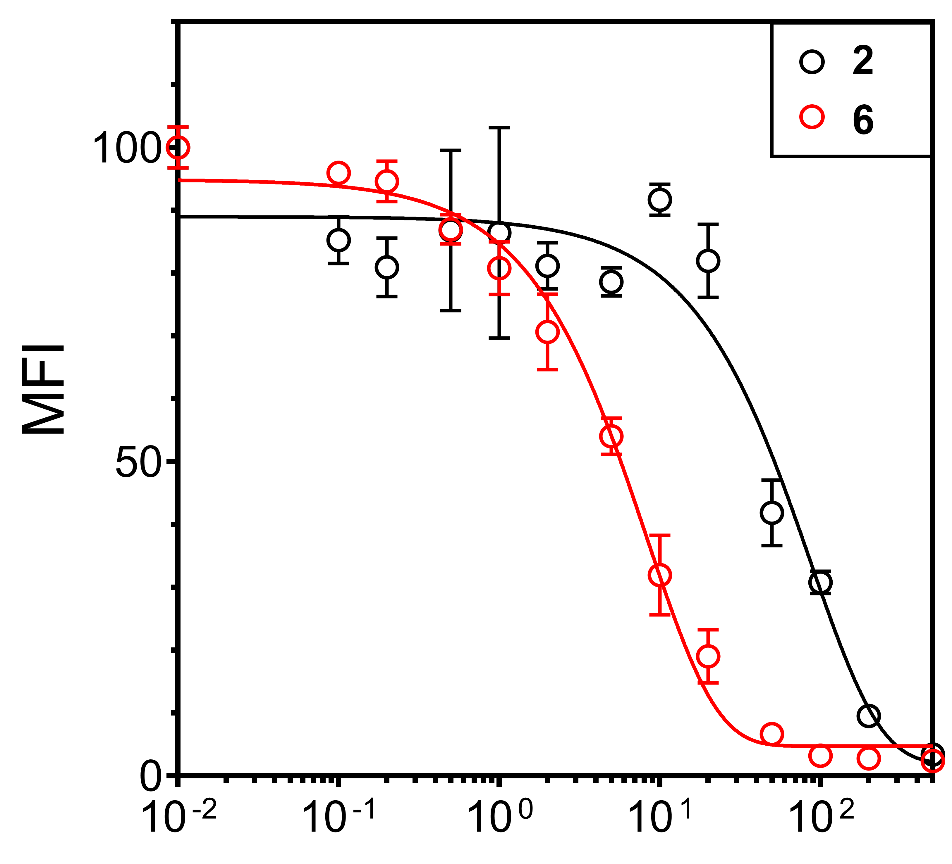

**Figure S10.** Flow cytometry analysis of *Msn* mc^2^ 155 incubated with 25 μM **TetD** then labeled with increasing concentrations of compound **2** or **6** followed by 50 μM **Fl-az** for 1h at 37 ℃. Data are represented as mean +/- SD (n = 3).

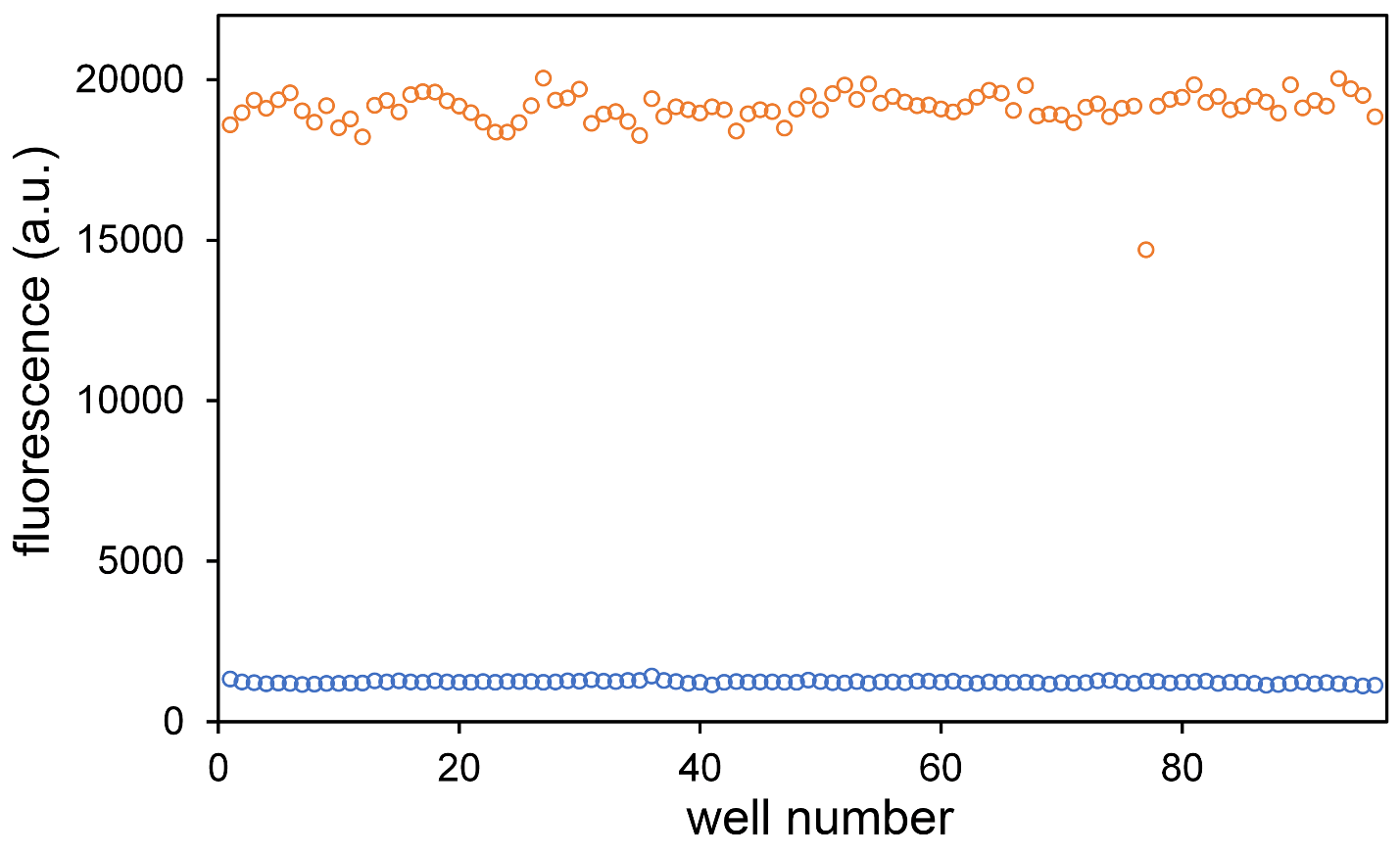

**Figure S11.** Mimicking of high throughput assays on 96-well plates. *Msn* mc^2^ 155 was incubated with 25 μM **TetD** were distributed to 2 96-well plates and treated with PBS or 50 μM compound **2** in the mini library for 2 h at 37 ℃. The cells were then labeled with 50 μM **Fl-Az** for 1h at 37 ℃. Data are represented as single dots representing 96 wells (n = 96). Blue, treatment of compound **2**, orange, treatment with PBS.

**Materials and Methods**

**General materials resources table.**

| **Reagents** | **Vendor Source** | **Catalog #** |
| --- | --- | --- |
| **Chemicals and synthesis materials** |  |  |
| α-N-Fmoc-amino acids | Chem Impex | Various |
| 2-Chlorotritul chloride resin | Chem Impex | 03498 |
| N, N’-diisopropylethylamine (DIEA) | Chem Impex | 00141 |
| Hexafluorophosphate Benzotriazole Tetramethyl Uronium (HBTU) | Chem Impex | 02011 |
| N,N-Dimethylformamide (DMF) ACS-grade | Millipore Sigma | 319937 |
| Dichloromethane ACS-grade | Millipore Sigma | D65100 |
| Methanol ACS-grade and HPLC grade | Millipore Sigma | 179337 |
| Acetonitrile ACS-grade and HPLC grade | Millipore Sigma |  |
| Piperidine | Millipore Sigma | 104094 |
| Trifluoroacetic acid (TFA) and HPLC grade | Millipore Sigma | 302031 |
| Dibenzocyclooctyne- DBCO-NHS ester | Broadpharm | BP-22231 |
| **Reagents and Media for bacterial assays** |  |  |
| Middlebrook 7H9 media | VWR | 90003-876 |
| Catalase from bovine liver | Millipore Sigma | C1345-1G |
| Dextrose | Chem Impex | 00805 |
| Bovine serum albumin fraction V | Millipore Sigma | 10735078001 |
| Glycerol | Millipore Sigma | G7893 |
| Tween 80 | VWR | 97061-674 |
| Formaldehyde | Millipore Sigma | 252549 |
| 5-carboxy fluorescein (**Fl-acid**) | Millipore Sigma | C0537 |
| 6-azido-fluorescein (**Fl-Az**) | Lumiprobe | D5130 |
| 6-DBCO-fluorescein (**Fl-DBCO**) | Lumiprobe | 451F0 |
| 6-azido-rhodamine 110 (**RoAz**) | Lumiprobe | D5230 |
| 3-azido-7-hydroxy coumarin (**CoAz**) | Millipore Sigma | 909513-5MG |
| Propidium iodide | Chem Impex | 00498 |
| Nile red | Chem Impex | 22855 |
| Amino terminated polystyrene beads | Spherotech | AP-30-10 |
| Azido molecule library | Various | Various |
| **Cell culture plates and assay plates** |  |  |
| 96-well clear conical bottom deep well plates | VWR | 76210-524 |
| 96-well clear untreated round bottom plates | VWR | 82050-622 |
| 96-well black half area black flat bottom plates | Millipore Sigma | CLS3694 |

**General Chemistry methods and Instruments**

Crude DBCO conjugates were purified by reverse phased preparative high performance liquid chromatography (RP-HPLC) equipped with Waters 1525 with 2489 UV/Visible Detector on a Phenomenex Luna 10 μm C8(2) 100 Å (250 x 21.2 mm) column using a 40 to 100% linear gradient of methanol in H_2_O/MeOH with 0.1% TFA at 10 mL/min. The HPLC fractions of the desired purified compounds were first concentrated under reduce pressure using rotary evaporator. The final concentrated aqueous solutions were lyophilized to dryness using Labconco Freezone 4.5L (-84^o^C) lyophilizer. The purity of the purified samples were analyzed with a Phenomenex Luna 5μm C8(2) on same RP-HPLC; gradient elution in H_2_O/CH_3_CN with 0.01% TFA at 1 mL/min. ^1^H and ^13^C-NMR Spectra for all conjugates for characterization were acquired on Varian 600MHz spectrophotometer. Residual solvent signal from DMSO-d6 referenced to tetramethylsilane (TMS) were used as reference standards for defining chemical shifts ^1^H or ^13^C spectra of compounds. Chemical shifts are reported in δ ppm and coupling constants (*J*) are reported in Hertz [Hz]. Deuterated solvents were used as received from Cambridge Isotopes. High resolution electrospray ionization mass spectrometry (HRMS, ESI/MS) analyses were obtained on an Agilent 6545B Q-TOF LC/MS equipped with 1260 infinity II LC system with auto sampler. Samples were dissolved in CH_3_CN and eluted with a CH_3_CN/H_2_O solution containing 0.1% formic acid. Concentrations of synthesized DBCO-conjugates for biological experiments were measured by absorbance at 309 nm (ε=12,000 cm^-1^M^-1^) by UV-visible spectrophotometer (Thermo Scientific Genesys-50).

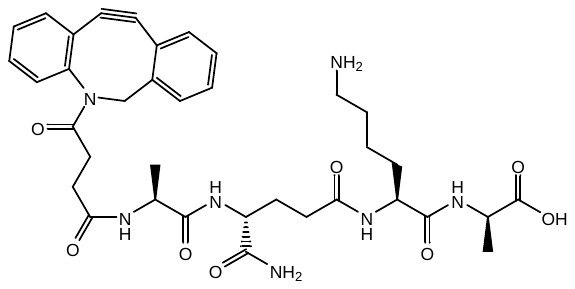
**Synthesis of TetD**

To a 25 mL peptide synthesis vessel with 100 mg 2-Chlorotrityl chloride resin (0.142 mmol) resuspended in 15 mL dry dichloromethane, was added Fmoc-D-alanine (49 mg, 1.1 eq, 0.16 mmol), and diisopropylethylamine (DIEA, 4.4 eq, 0.11 mL, 0.62 mmol). The resin was shaken for 1 hour at room temperature and washed with methanol and dichloromethane (3 times and 15 mL each). Fmoc protecting group was removed with 6M piperazine in N, N-Dimethylformamide (DMF, 15 mL) for 30 min at room temperature and washed as before. Fmoc-L-Lys (Boc)-OH (3.0 eq, 0.20 g, 0.43 mmol), HBTU (3.0 eq, 0.16 g, 0.43 mmol), and DIEA (6.0 eq, 0.15 mL, 0.85 mmol) in DMF (15 mL) was added to the vessel and shaken for 2 h at room temperature. The Fmoc deprotection and coupling procedure was repeated using the same equivalent with Fmoc-D-glutamic acid α-amide and Fmoc-L-alanine.

DBCO was coupled on the N term of the tetra peptide on resin. 25-30 mg DBCO-NHS was dissolved in 1 mL dry DMF and added to the 25 mL peptide synthesis vessel with 100 mg equivalent 2-Chlorotrityl chloride resin with tetra peptide resuspended in 2 mL DMF. The resin was shaken overnight at room temperature and washed with methanol and dichloromethane (3 times and 15 mL each). The resin was then added 20% trifluoroacetic acid (TFA) in dichloromethane after wash and shaken in room temperature for 1 h. The liquid phase was filtered and concentrated with nitrogen flow and added icy ether to precipitate the peptide. The ether layer was decanted, and the resulting solid was washed with icy ether and air dried. The crude material was purified with reverse phased high performance liquid chromatography (RP-HPLC). The purified sample was analyzed for purity using a Waters 1525 with a Phenomenex analytical column (R.t. 17.7 min). Partial ^1^H-NMR (DMSO-d_6_) mixture of rotamers δ 1.12 (d, *J*= 6Hz, 3H, CH_3_ minor), 1.16 (d, *J*=6Hz, 3H, CH_3_ major), 1.24-1.25 (dd, 6H, CH_3_), 1.28 (m, 2H, CH_2_), 1.50 (m, 4H), 1.65 (m, 2H), 1.78 (m, 2H), 1.91-2.12 (m, 6H), 2.24(q, *J*=12Hz, 16Hz, 1H), 2.30 (q, *J*=12Hz, 16Hz, 1H), 2.61 (m, 2H), 2.73 (m, 4H), 3.61 (dd, *J*=12Hz, 18Hz, 2H), 4.05-4.15 (m, 4H), 4.20 (m, 2H), 4.26 (m, 2H), 5.04 (t, *J*=12Hz, 4H), 7.08 (d, *J*=12Hz, 2H), 7.20 (d, *J*=6Hz, 2H), 7.30 (d, *J*=12Hz, 2H), 7.34 (t, *J*=12Hz, 2H), 7.37 (t, *J*=12Hz, 2H), 7.46-7.51 (m, 6H), 7.62-7.67 (m, 8H), 7.86 (d, *J*=6Hz, 1H), 7.90 (t, *J*=12Hz, 2H), 7.95 (d, *J*=6Hz, 1H), 8.05 (d, *J*=6Hz, 2H), 8.13 (d, *J*=6Hz, 2H). Partial ^13^C-NMR (DMSO-d_6_) major rotamer δ 17.57, 17.63, 22.24, 26.63, 26.65, 27.76, 29.52, 30.02, 30.23, 31.65, 40.06, 47.41, 48.64, 51.96, 52.03, 54.89, 108.07, 114.25, 114.94, 116.89, 121.41, 122.49, 125.15, 126.81, 127.71, 128.05, 129.61, 132.44, 140.72, 148.40, 151.45, 151.55, 158.34, 171.16, 171.28, 171.39, 171.48, 171.59, 171.67, 172.36, 173.27, 173.92. HRMS [QTOF-MS]: calculated for C36H46N7O8, 704.3402, found: (M+H)^+^ 704.3404.

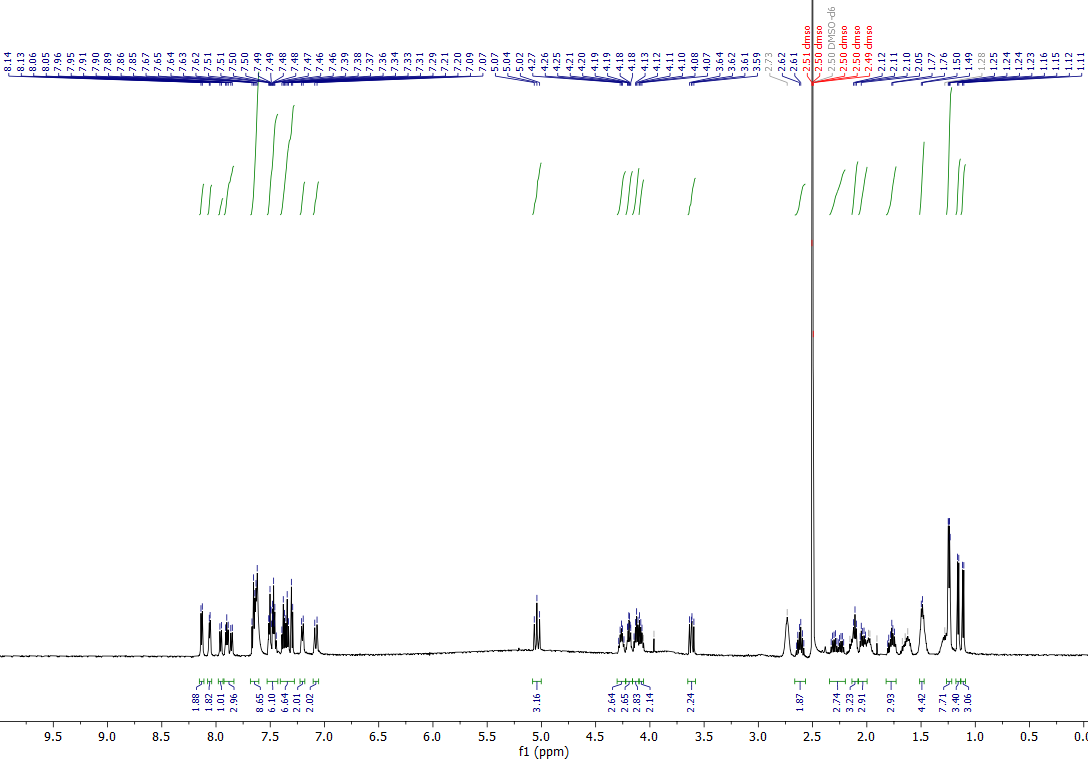

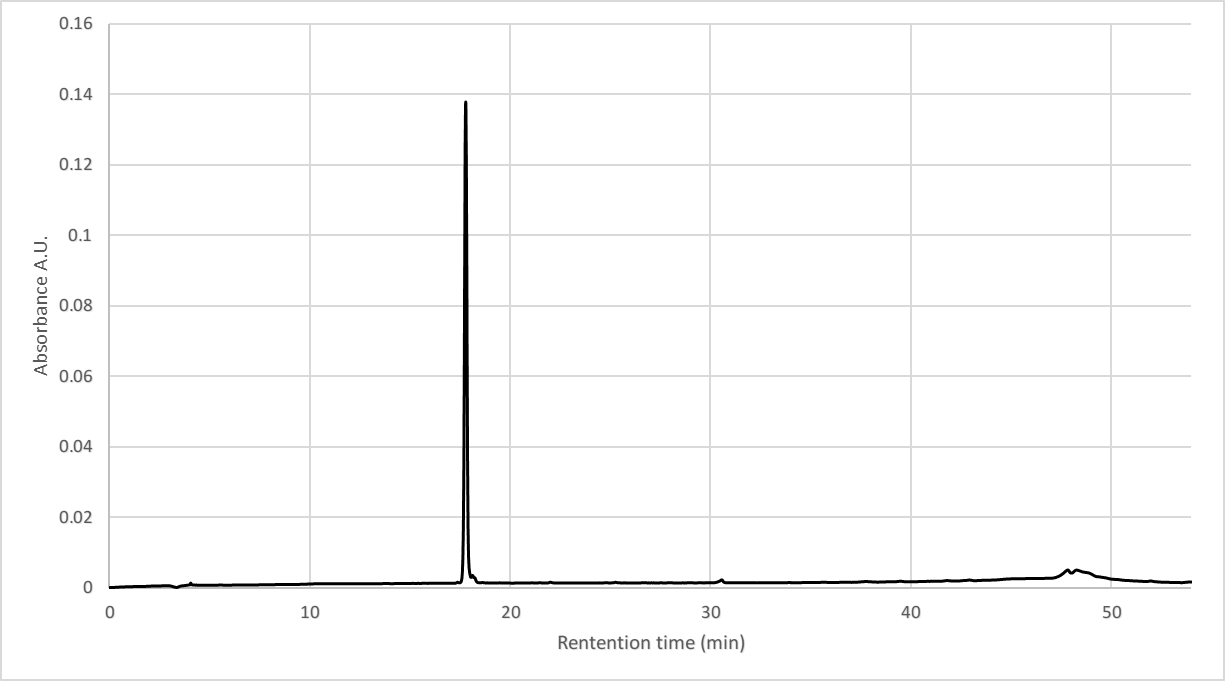

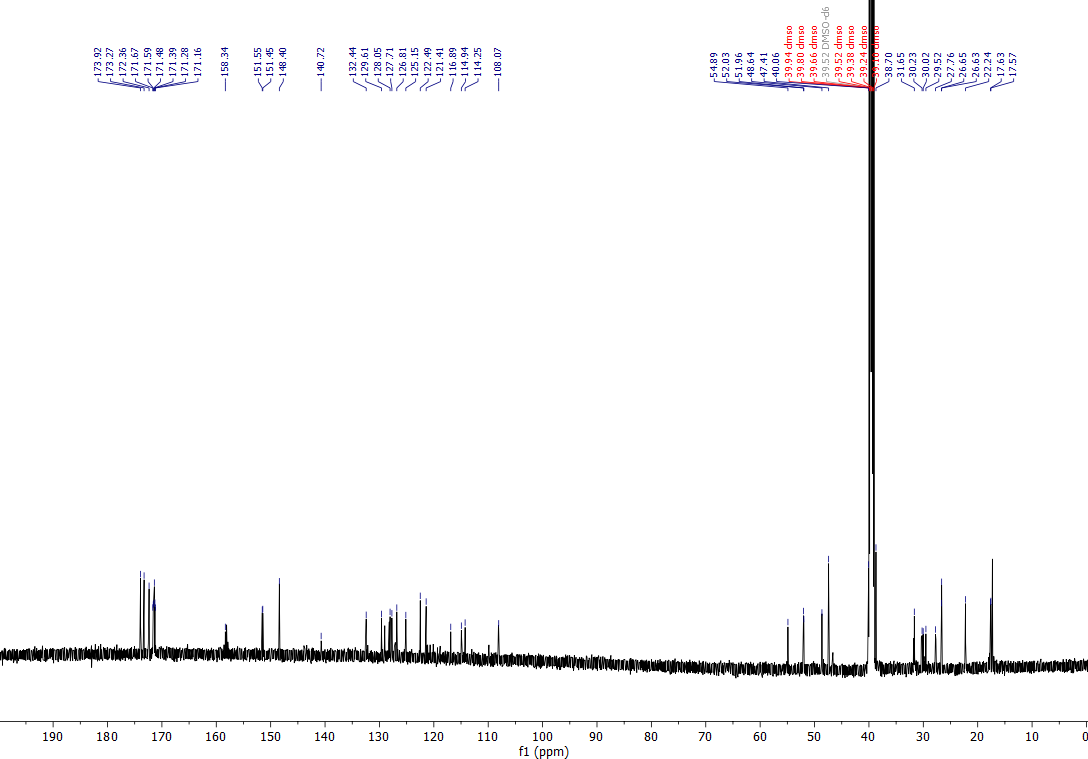

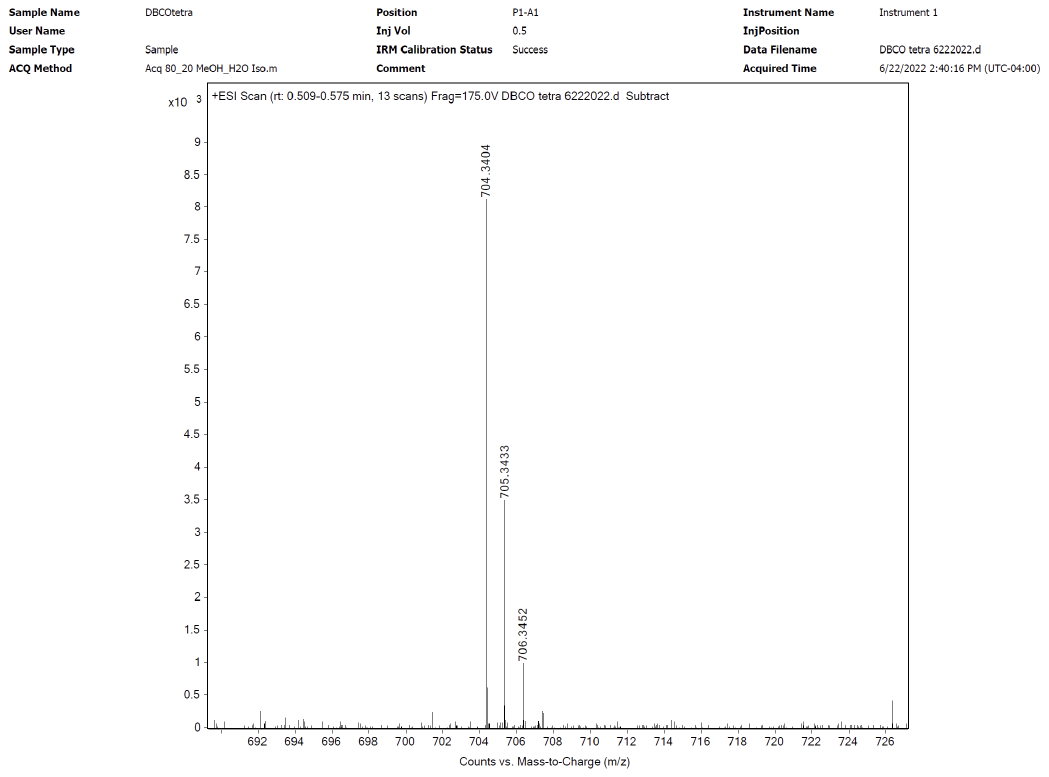

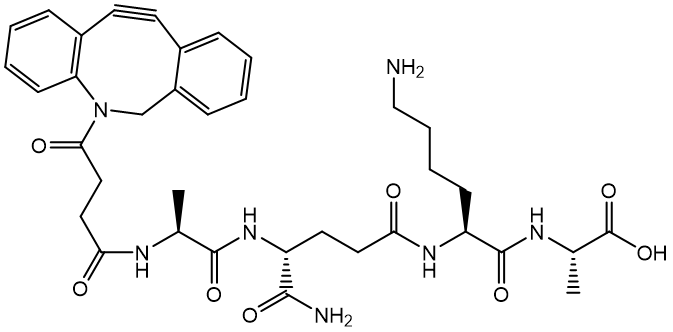
**Synthesis of TetD(L-control)**

To a 25 mL peptide synthesis vessel with 100 mg 2-Chlorotrityl chloride resin (0.142 mmol) resuspended in 15 mL dry dichloromethane, was added Fmoc-L-alanine (49 mg, 1.1 eq, 0.16 mmol), and diisopropylethylamine (DIEA, 4.4 eq, 0.11 mL, 0.62 mmol). The resin was shaken for 1 hour at room temperature and washed with methanol and dichloromethane (3 times and 15 mL each). Fmoc protecting group was removed with 6M piperazine in N, N-Dimethylformamide (DMF, 15 mL) for 30 min at room temperature and washed as before. Fmoc-L-Lys (Boc)-OH (3.0 eq, 0.20 g, 0.43 mmol), HBTU (3.0 eq, 0.16 g, 0.43 mmol), and DIEA (6.0 eq, 0.15 mL, 0.85 mmol) in DMF (15 mL) was added to the vessel and shaken for 2 h at room temperature. The Fmoc deprotection and coupling procedure was repeated using the same equivalent with Fmoc-D-glutamic acid α-amide and Fmoc-L-alanine.

DBCO was coupled on the N term of the tetra peptide on resin. 25-30 mg DBCO-NHS was dissolved in 1 mL dry DMF and added to the 25 mL peptide synthesis vessel with 100 mg equivalent 2-Chlorotrityl chloride resin with tetra peptide resuspended in 2 mL DMF. The resin was shaken overnight at room temperature and washed with methanol and dichloromethane (3 times and 15 mL each). The resin was then added 20% trifluoroacetic acid (TFA) in dichloromethane after wash and shaken in room temperature for 1 h. The liquid phase was filtered and concentrated with nitrogen flow and added icy ether to precipitate the peptide. The ether layer was decanted, and the resulting solid was washed with cold ether and air dried. The crude material was further purified with reverse phase high performance liquid chromatography (RP-HPLC). The purified sample was analyzed for purity using a Waters 1525 with a Phenomenex analytical column (R.t. 18.0 min). Partial ^1^H-NMR (DMSO-d_6_) mixture of rotamers δ 1.10 (d, *J*= 6Hz, 3H), 1.15 (d, *J*=6Hz, 3H), 1.24 (d, *J*=6Hz, 3H, CH_3_), 1.27 (d, *J*=6Hz, 3H, CH_3_), 1.31 (m, 2H, CH_2_), 1.44-1.55 (m, 4H), 1.63 (m, 2H), 1.75 (m, 2H), 1.96-2.15 (m, 6H), 2.22(q, *J*=12Hz, 16Hz, 1H), 2.30 (q, *J*=12Hz, 16Hz, 1H), 2.59 (m, 2H), 2.74 (m, 4H), 3.61 (t, *J*=12Hz, 2H), 4.08-4.20 (m, 6H), 4.26 (m, 2H), 5.04 (t, *J*=12Hz, 4H), 7.08 (d, *J*=12Hz, 2H), 7.19 (d, *J*=6Hz, 2H), 7.30 (d, *J*=12Hz, 2H), 7.34 (t, *J*=12Hz, 2H), 7.38 (t, *J*=12Hz, 2H), 7.46-7.51 (m, 6H), 7.62-7.67 (m, 8H), 7.84 (d, *J*=12Hz, 1H), 7.93 (t, *J*=12Hz, 2H), 7.95 (d, *J*=6Hz, 1H), 8.06 (t, *J*=6Hz, 2H), 8.17 (t, *J*=6Hz, 2H). HRMS [QTOF-MS]: calculated for C36H46N7O8, 704.3402, found: (M+H)^+^ 704.3406.

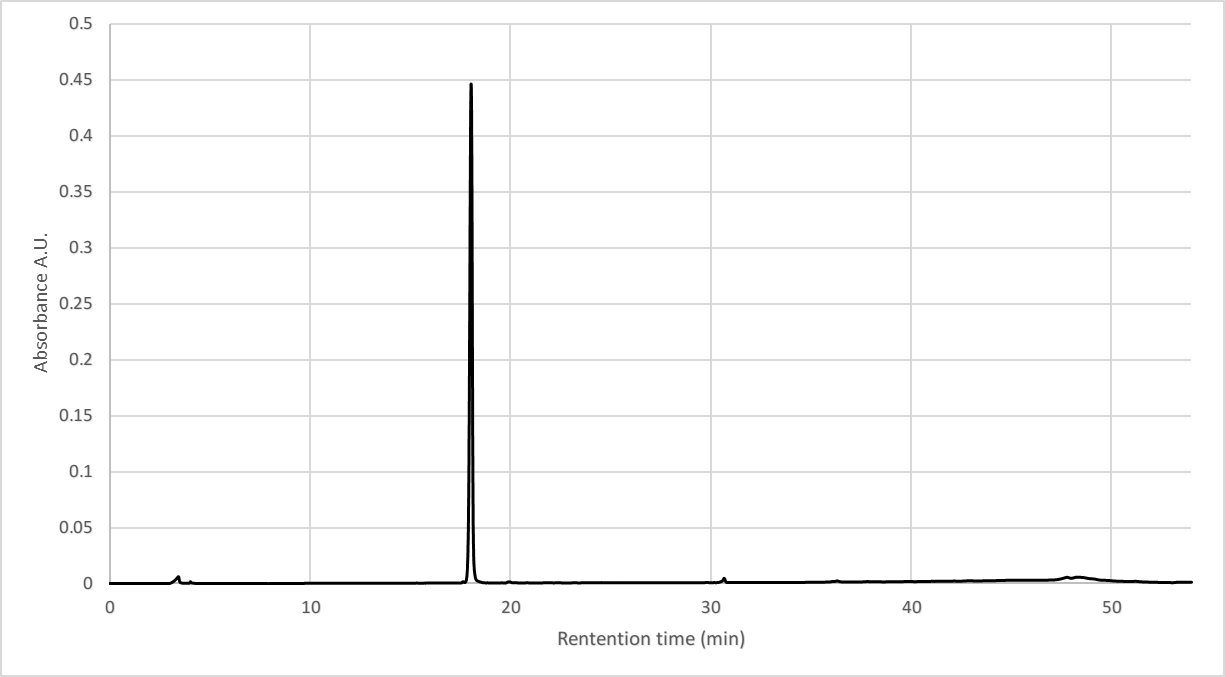

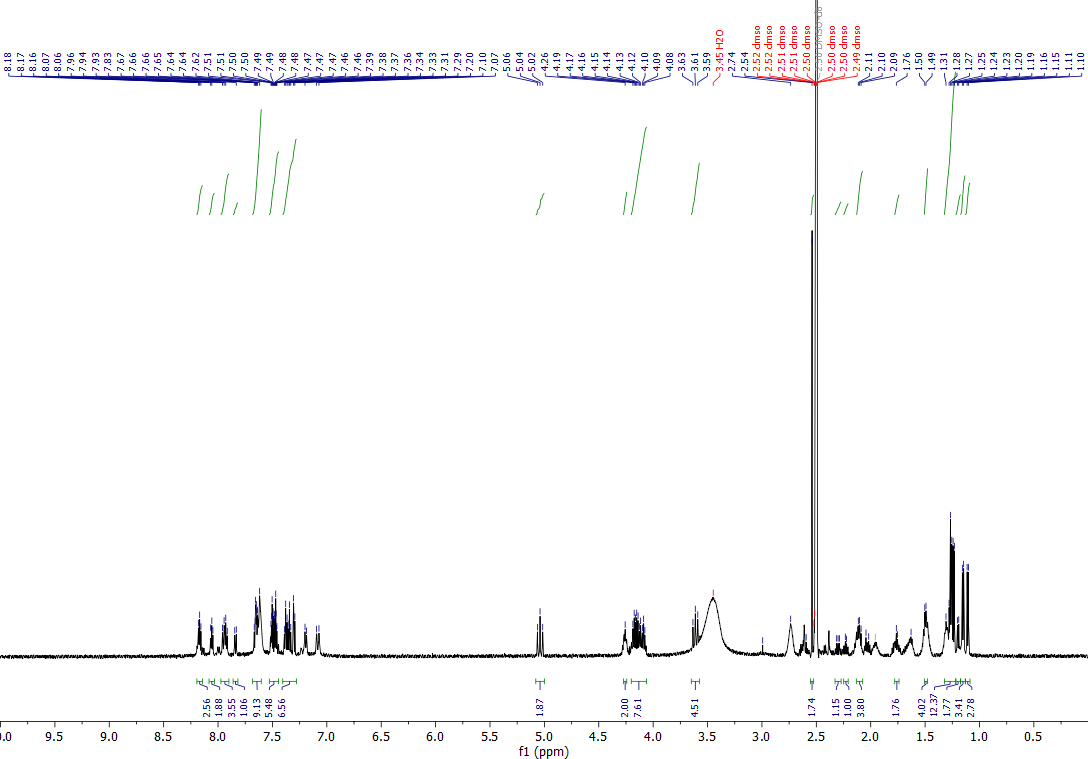

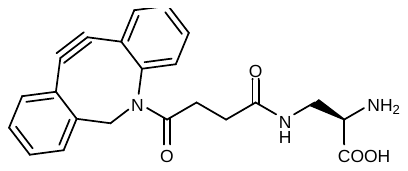

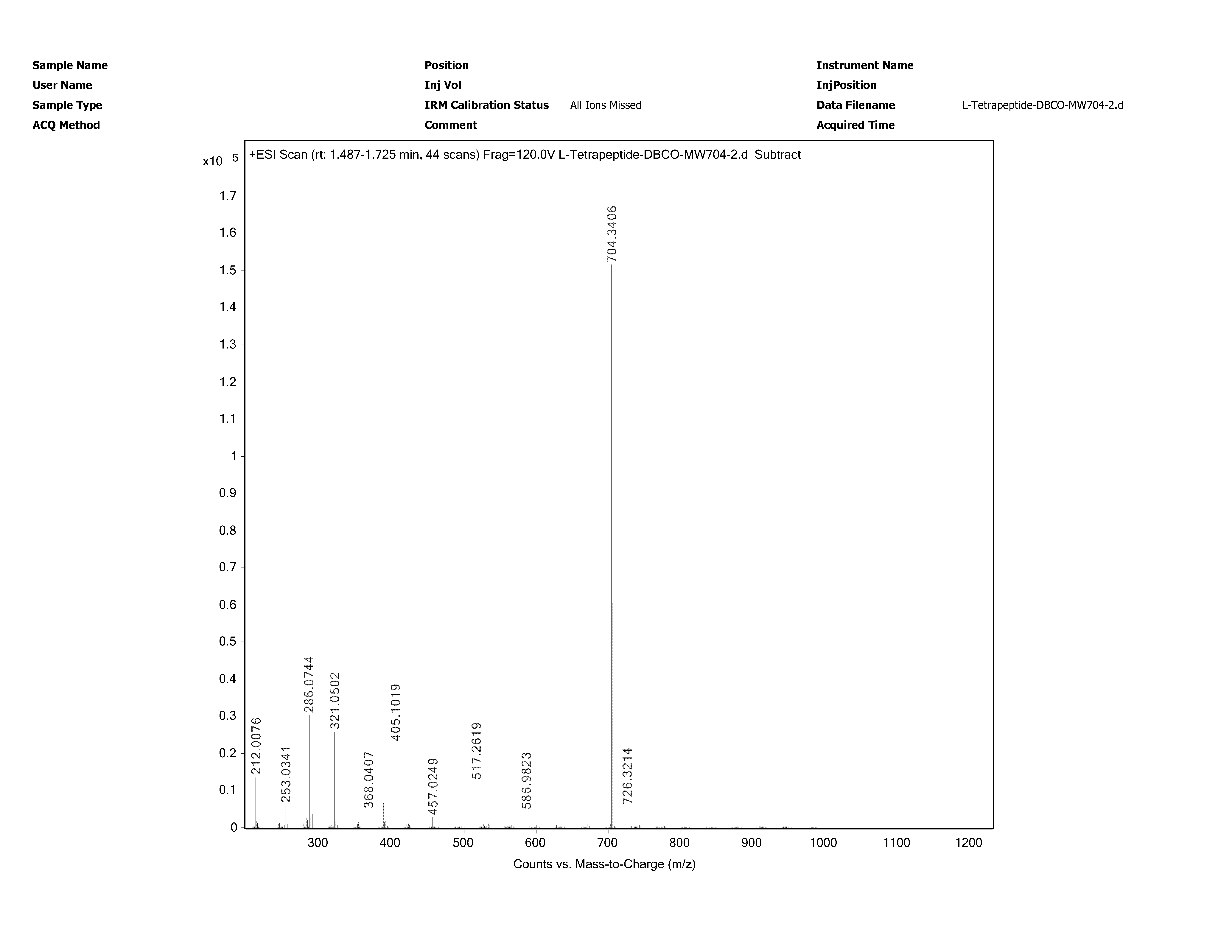
**Synthesis of D-DapD**

To a 25 mL peptide synthesis vessel with 100 mg 2-Chlorotrityl chloride resin (0.142 mmol) resuspended in 15 mL dry dichloromethane, was added Nα-Boc-Nβ-Fmoc-D-2,3-diaminopropionic acid (D-Dap, 67 mg, 1.1 eq, 0.16 mmol), and DIEA (4.4 eq, 0.11 mL, 0.62 mmol). The resin was shaken for 1 hour at room temperature and washed with methanol and dichloromethane (3 times and 15 mL each). Fmoc protecting group was removed with 6M piperazine in N, N-Dimethylformamide (DMF, 15 mL) for 30 min at room temperature and washed as before.

DBCO was coupled on the side chain of D-Dap on resin. 25-30 mg DBCO-NHS was dissolved in 1 mL dry DMF and added to the 25 mL peptide synthesis vessel with 100 mg equivalent 2-Chlorotrityl chloride resin with D-Dap resuspended in 2 mL DMF. The resin was shaken overnight at room temperature and washed with methanol and dichloromethane (3 times and 15 mL each). The resin was then added 20% trifluoroacetic acid (TFA) in dichloromethane after wash and shaken in room temperature for 1 h. The liquid phase was filtered and concentrated with nitrogen flow and added icy ether to precipitate the peptide. The ether layer was decanted, and the resulting solid was washed with icy ether and air dried. The crude material was purified with reverse phased high performance liquid chromatography (RP-HPLC). The purified sample was analyzed for purity using a Waters 1525 with a Phenomenex analytical column (R.t. 19.7 min). ^1^H-NMR (DMSO-d_6_) mixture of rotamers δ 1.79-1.86 (m, 2H), 2.04-2.10 (m, 2H), 2.22-2.32 (m, 2H), 3.27-3.32 (m, 2H), 3.53-3.57 (m, 2H), 3.63 (d, *J*=12Hz, 2H), 3.90 (brs, 2H), 5.03 (d, *J*=18Hz, 2H), 7.30 (d, *J*=6Hz, 2H), 7.36 (t, *J*=6Hz, 2H), 7.40 (dt, *J*=2Hz, 6Hz, 2H), 7.45-7.52 (m, 6H), 7.63 (d, *J*=12Hz, 2H), 7.66 (dt, *J*=2Hz, 6Hz, 2H), 8.00 (t, *J*=6Hz, 2H), 8.02 (t, *J*=6Hz, 2H), 8.10 (brs). ^13^C-NMR (DMSO-d_6_) mixture of rotamers δ 29.49, 29.57, 30.31, 38.55, 52.33, 54.94, 108.12, 114.27, 121.45, 122.49, 125.18, 126.85, 127.72, 128.05, 128.23, 128.97, 129.58, 132.45, 148.40, 151.52, 169.16, 171.11, 171.14, 172.70. HRMS [QTOF-MS]: calculated for C22H22N3O4, 392.1605, found: (M+H)^+^ 392.1607.

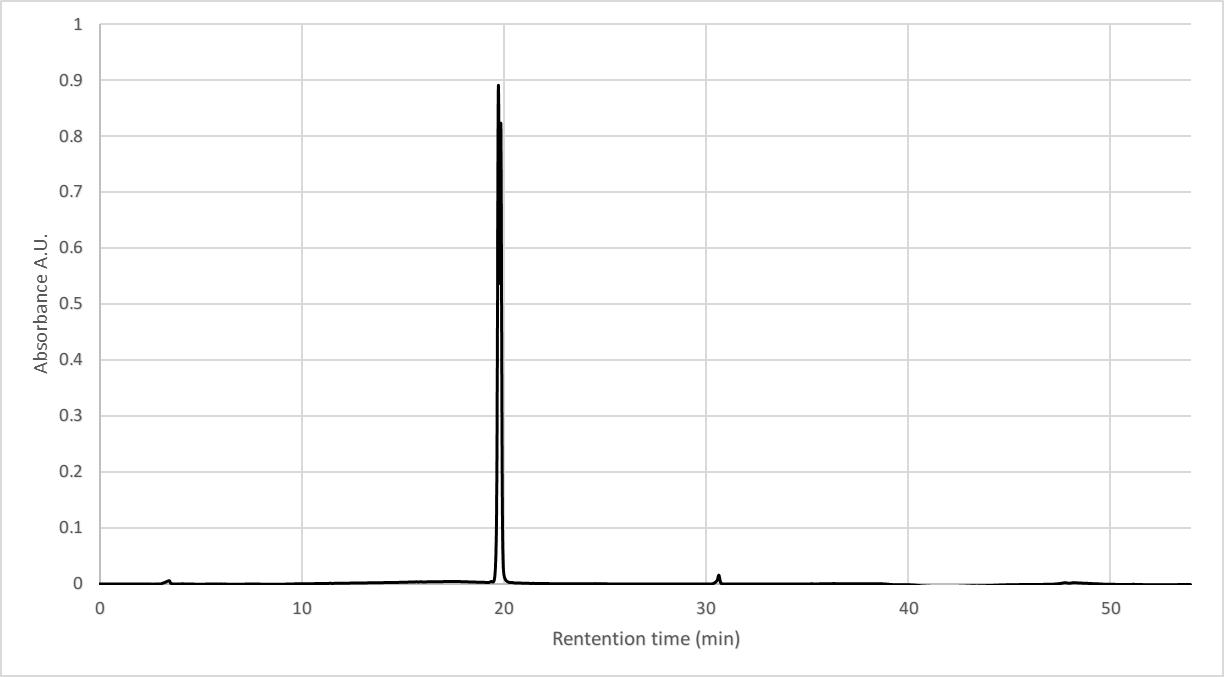

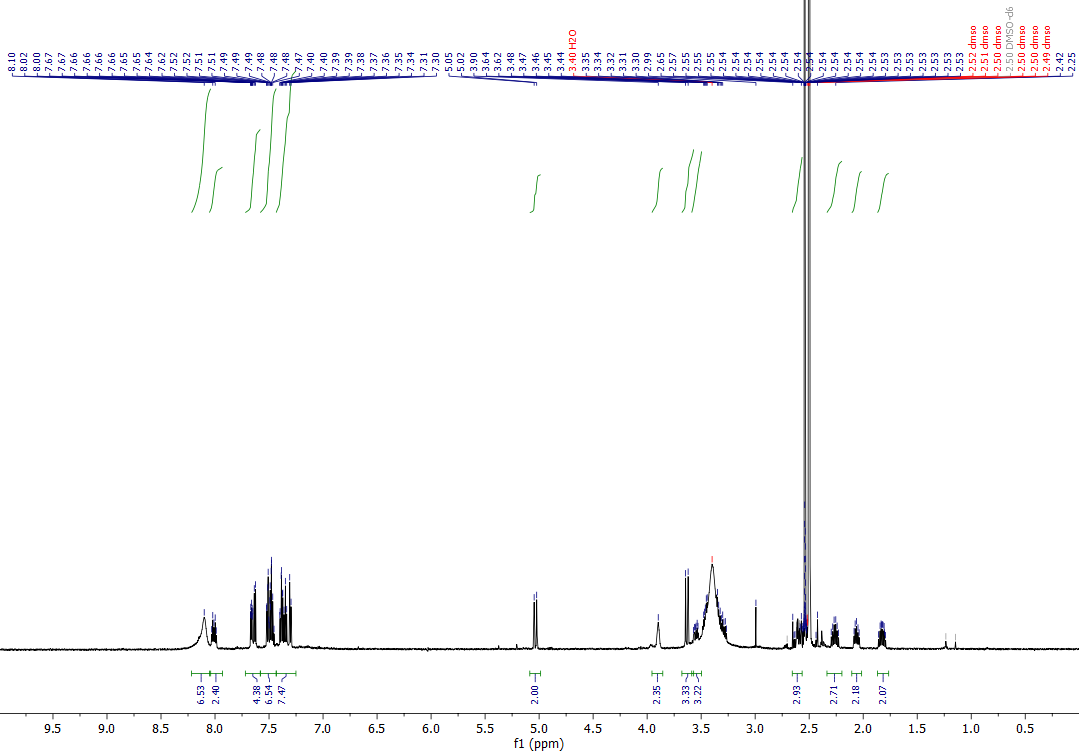

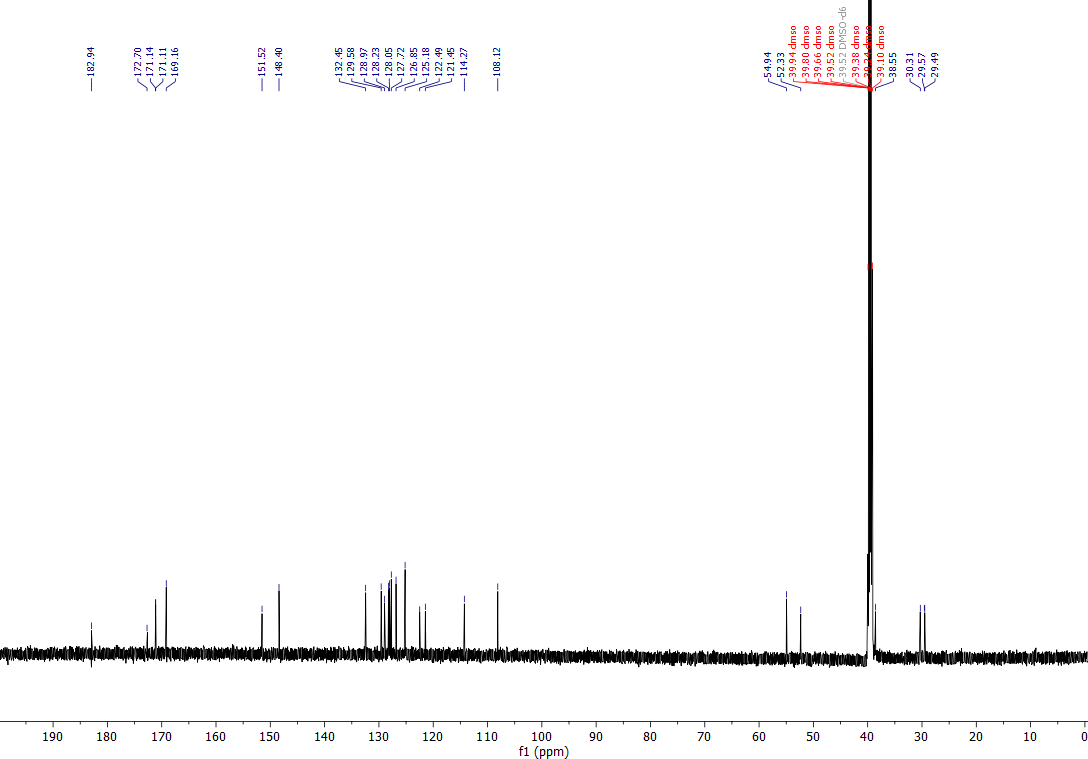

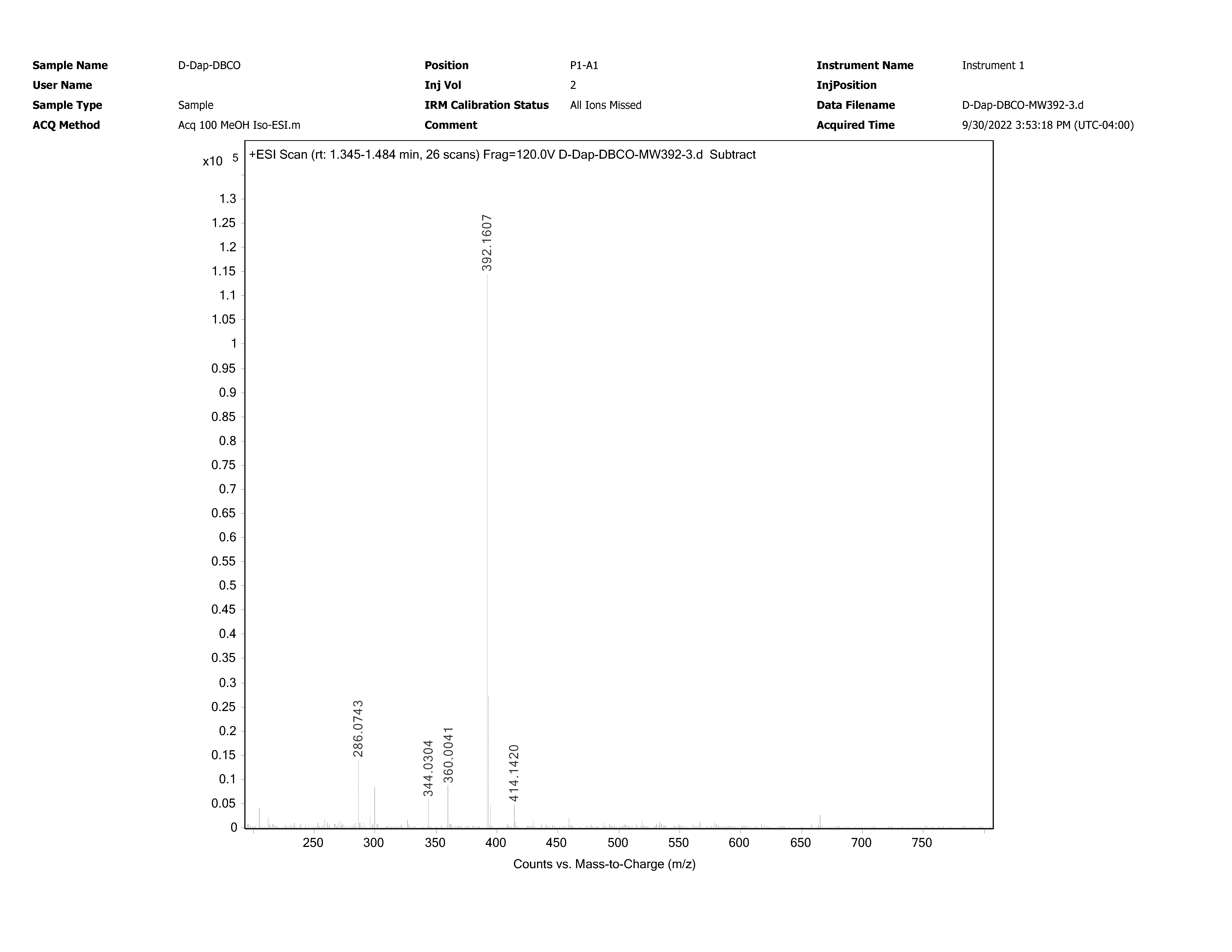

**General Cell Culture.** *Mycobacterium smegmatis* strains mc^2^ 155, ATCC 14468, and PM2750 ∆5 were grown in 7H9 media with 0.5% glycerol, 0.05% tween 80, and 1x ADC enrichment (10x ADC, 5 g bovine serum albumin, 2 g dextrose, 3 mg catalase in 100 mL). *M. smegmatis* mc^2^ 1255 was grown in the same media with 50 μg/mL streptomycin sulfate. Glycerol stocks were made using stationary phase cells in 30% glycerol and aliquots were stored in - 80 ℃.

**General method for bacterial peptidoglycan modification**. *M. smegmatis* strains were inoculated from the glycerol stock to the according media and grown for 24 hours until 0.5-0.6 OD, tetrapeptide probes were added to the media and the cells were grown overnight to achieve stationary phase. The cells were harvested the next day and spun down for 2 min at 3000 g, washed with phosphate buffered saline with tween 80 (PBST, phosphate buffered saline (PBS) with 0.05% tween 80) two times and resuspended in PBST to yield DBCO-modified *M. smegmatis* cells for further labeling and other experiments.

**Comparison with stereo control and DBCO-modified single amino acid.** 30 μL *M. smegmatis* mc^2^ 155 was inoculated from stationary phase in 1:100 dilution to 3 mL fresh 7H9 media with ADC. 25 μM **D-DapD**, **TetD** and **TetD** (L-control) were added to the culture tubes respectively, and the cells were grown for 36-40 hours until stationary phase. The cells were harvested and spun down for 2 min at 3000 g, washed with 3 mL PBST two times and resuspended in 3 mL PBST. To a 96-well plate added 90 μL cells pre well with 10 μL 500 μM **Fl-Az** (final 50 μM) in triplicate. The plate was incubated in 37 ℃ for 1 h and spun down for 2 min at 2700 g. The supernatant was decanted, and the pellets were washed with 100 μL PBST for 2 times and fixed with 100 μL 4% formaldehyde for 15 min. The samples were then analyzed by Attune^TM^ NxT Acoustic Focusing Cytometer.

**Confirm for click chemistry on PG.** 30 μL *M. smegmatis* mc^2^ 155 was inoculated from stationary phase in 1:100 dilution to 3 mL fresh 7H9 media with ADC. 25 μM **TetD** and **TetFl** were added to the culture tubes respectively, and the cells were grown for 36-40 hours until stationary phase. The cells were harvested and spun down for 2 min at 3000 g, washed with 3 mL PBST two times and resuspended in 3 mL PBST. To a 96-well plate added 90 μL cells pre well with 10 μL 500 μM **Fl-Az** or **Fl-acid** (50 μM final) accordingly in triplicate. The plate was incubated in 37 ℃ for 1 h and spun down for 2 min at 2700 g. The supernatant was decanted, and the pellets were washed with 100 μL PBST for 2 times and fixed with 100 μL 4% formaldehyde for 15 min. The samples were then analyzed by Attune^TM^ NxT Acoustic Focusing Cytometer.

**Concentration dependency on TetD.** 30 μL *M. smegmatis* mc^2^ 155 was inoculated from stationary phase in 1:100 dilution to 3 mL fresh 7H9 media with ADC. 5 μM, 10 μM or 25 μM **TetD** was added to the culture tubes respectively, and the cells were grown for 36-40 hours until stationary phase. The cells were harvested and spun down for 2 min at 3000 g, washed with 3 mL PBST two times and resuspended in 3 mL PBST. To a 96-well plate added 90 μL cells pre well with 10 μL 500 μM **Fl-Az** (50 μM final) in triplicate. The plate was incubated in 37 ℃ for 1 h and spun down for 2 min at 2700 g. The supernatant was decanted, and the pellets were washed with 100 μL PBST for 2 times and fixed with 100 μL 4% formaldehyde for 15 min. The samples were then analyzed by Attune^TM^ NxT Acoustic Focusing Cytometer.

**Concentration and temporal dependency of Fl-Az.** 30 μL *M. smegmatis* mc^2^ 155 was inoculated from stationary phase in 1:100 dilution to 3 mL fresh 7H9 media with ADC. 25 μM **TetD** was added to the culture tubes, and the cells were grown for 36-40 hours until stationary phase. The cells were harvested and spun down for 2 min at 3000 g, washed with 3 mL PBST two times and resuspended in 3 mL PBST. To a 96-well plate added 40 μL cells pre well with 60 μL PBST with different amount of **Fl-Az** to make up the concentrations described in the main text, 9 wells each. The plate was incubated in 37 ℃, the cells from 3 wells in each concentration group were taken out at 30 min, 1 h, or 2h and spun down for 2 min at 2700 g. The supernatant was decanted, and the pellets were washed with 100 μL PBST for 2 times and fixed with 100 μL 4% formaldehyde for 15 min. The samples were then analyzed by Attune^TM^ NxT Acoustic Focusing Cytometer.

**Dye comparison.** 30 μL *M. smegmatis* mc^2^ 155 was inoculated from stationary phase in 1:100 dilution to 3 mL fresh 7H9 media with ADC. 25 μM **TetD** was added to the culture tubes, and the cells were grown for 36-40 hours until stationary phase. The cells were harvested and spun down for 2 min at 3000 g, washed with 3 mL PBST two times and resuspended in 3 mL PBST. To a 96-well plate added 90 μL cells pre well with 10 μL 500 μM **Fl-Az**, **RoAz** or **CoAz**, respectively (50 μM final), in triplicate. The plate was incubated in 37 ℃ for 1 h and spun down for 2 min at 2700 g. The supernatant was decanted, and the pellets were washed with 100 μL PBST for 2 times and fixed with 100 μL 4% formaldehyde for 15 min. The samples were then analyzed by Attune^TM^ NxT Acoustic Focusing Cytometer. BL1 channel was used for **Fl-Az** and **RoAz**, VL1 channel was used for **CoAz**.

**L, D-transpeptidase knock down strain.** *M. smegmatis* mc^2^ 155 and PM2750 ∆5 were inoculated from the glycerol stock to 3 mL of the according media and grown for 48 hours to reach stationary phase, then 3 μL from each stationary culture was inoculated to 3 mL fresh 7H9 media with ADC and grown for 24 h until 0.5-0.6 OD.25 μM **TetD** was added to the media, and the cells were grown overnight to achieve stationary phase. The cells were harvested and spun down for 2 min at 3000 g, washed with 3 mL PBST two times and resuspended in 3 mL PBST. To a 96-well plate added 90 μL cells pre well with 10 μL 500 μM **Fl-Az** (50 μM final) accordingly in triplicate. The plate was incubated in 37 ℃ for 1 h and spun down for 2 min at 2700 g. The supernatant was decanted, and the pellets were washed with 100 μL PBST for 2 times and fixed with 100 μL 4% formaldehyde for 15 min. The samples were then analyzed by Attune^TM^ NxT Acoustic Focusing Cytometer.

**L, D-transpeptidase inhibition with meropenem.** *M. smegmatis* mc^2^ 155 was inoculated from the glycerol stock by 1 to 1000 dilution to 3 mL fresh 7H9 media with ADC and grown for 24 hours until 0.5-0.6 OD. 25 μM **TetD** was added to the media, and the cells were aliquoted to a 96-well culture plate, each well consisting of 160 μL cell media and 40 μL PBS with different amount of meropenem to make op the concentrations described in the main text, in triplicate respectively. The cells were then grown overnight to achieve stationary phase. 100 μL cells were transferred to a 96-well plate and spun down for 2 min at 2700 g, washed with 100 μL PBST each well two times and resuspended in 100 μL 50 μM **Fl-Az** in PBST. The plate was incubated in 37 ℃ for 1 h and spun down for 2 min at 2700 g. The supernatant was decanted, and the pellets were washed with 100 μL PBST for 2 times and fixed with 100 μL 4% formaldehyde for 15 min. The samples were then analyzed by AttuneTM NxT Acoustic Focusing Cytometer.

**Competition with test molecules.** To a 96-well plate added 90 μL DBCO-modified *M. smegmatis* mc^2^ 155 cells pre well with 10 μL 500 μM test molecules, (50 μM final concentration) in triplicate respectively. The plate was incubated in 37 ℃ for 2 h. The cells were spun down for 2 min at 2700 g. The supernatant was decanted, and 100 μL 50 μM **Fl-Az** was then added to each well, followed by incubation in 37 ℃ for 1 h. The cells were then spun down again, and the pellets were washed with 100 μL PBST for 2 times and fixed with 100 μL 4% formaldehyde for 15 min. The samples were then analyzed by Attune^TM^ NxT Acoustic Focusing Cytometer.

**EC50 curves.** To a 96-well plate added 50 μL DBCO-modified *M. smegmatis* mc^2^ 155 cells pre well with 50 μL PBS containing different amount of test molecule to make up the described concentrations in the main text, in triplicate respectively. The plate was incubated in 37 ℃ for 2 h. The cells were spun down for 2 min at 2700 g. The supernatant was decanted, and 100 μL 50 μM **Fl-Az** was then added to each well, followed by incubation in 37 ℃ for 1 h. The cells were then spun down again, and the pellets were washed with 100 μL PBST for 2 times and fixed with 100 μL 4% formaldehyde for 15 min. The samples were then analyzed by Attune^TM^ NxT Acoustic Focusing Cytometer.

**Screen of the 12-plate library.** *M. smegmatis* mc^2^ 155 was inoculated from the glycerol stock by 1 to 1000 dilution to 50 mL fresh 7H9 media with ADC in 250 mL Erlenmeyer flasks each day and grown for 24 hours until 0.5-0.6 OD. 25 μM **TetD** were added to the media, and the cells were grown overnight to achieve stationary phase. The cells were harvested and spun down for 10 min at 3000 g, washed with 50 mL PBST two times and resuspended in 50 mL PBST. Each day, to 4 96-well plates added 90 μL cells with 10 μL 500 μM of each molecule each well (50 μM final). The plates were incubated in 37 ℃ for 2 h. The cells were spun down for 2 min at 2700 g. The supernatant was decanted, and 100 μL 50 μM **Fl-Az** was then added to each well, followed by incubation in 37 ℃ for 1 h. The cells were then spun down again, and the pellets were washed with 100 μL PBST for 2 times and fixed with 100 μL 4% formaldehyde for 15 min. The samples were then analyzed by Attune^TM^ NxT Acoustic Focusing Cytometer.

**Ethidium bromide and Nile red whole-cell accumulation assay.** To a Costar 96-well half area black opaque flat bottom plate added 95 μL DBCO-modified *M. smegmatis* cells each well with 5 μL 100 μM ethidium bromide (5 μM final) and 5 μL 200 μM Nile red (10 μM final) in triplicates, respectively. The fluorescent intensity at different time points were taken by a Synergy H1 microplate reader for 90 min with 3 min intervals and continuously orbital shaking. Wavelengths ethidium bromide, excitation 530 nm, emission 590 nm; Nile red, excitation 540 nm, emission 630 nm.

**Isolation of peptidoglycan and muropeptides.** *M. smegmatis* mc^2^ 155 was inoculated from the glycerol stock by 1 to 1000 dilution to 3 mL fresh 7H9 media with ADC and grown for 24 h to reach 0.5-0.6 OD. 25 mM **TetD**, **TetFl**, and **2-TreAz** were added to the culture tubes respectively, and the cells were grown overnight to achieve stationary phase. The cells were harvested and spun down for 2 min at 3000 g, washed with 4 mL PBST two times and resuspended in 4 mL PBST. 180 μL cells with DBCO-modification were aliquoted to a 96-well culture plate with 20 μL 500 μM **Fl-Az** (50 μM final) and incubated in 37 ℃ for 1 h. 180 μL cells labeled with 2-TreAz were added to the same 96-well culture plate with 20 μL 500 μM **Fl-DBCO** (50 μM final) and incubated in 37 ℃ for 1h. Blank cells and the **TetFl** labeled cells were aliquoted the same way with PBST to make up the volume. After incubation, the cells were spun down at 2700 g for 10 min and washed with 200 μL PBST two times. The cell pellets were resuspended in 200 μL 10 mM NH_4_HCO_3_ with protease inhibitor and bath sonicated for 30 min. 10 μg/mL DNase and RNase were added to each well directly after sonication and the plate was placed in 4 ℃ for 1 h. The cell wall-enriched fraction was collected by centrifugation at 2700 g for 10 min. The pellets were then treated with 200 μL PBS with 2% sodium dodecyl sulfate (SDS) each well and incubated at 50 ℃ for 1 h with shaking at 250 rpm. The suspension was spun down at 2700 g for 10 min. This treatment was repeated two times. Then the resulting pellet was resuspended in 200 μL PBS with 1% SDS and 0.1 mg/ml proteinase K each well, incubated at 37 ℃ for 1h with shaking at 250 rpm. The suspension was then heated with boiling water for 1 h and then spun down at 2700 g for 10 min. The supernatant was discarded and the 1% SDS extraction step was repeated 2 times. The pellet was then washed twice with PBS and 4 times with deionized water to give mycolyl-arabinogalactan-peptidoglycan Complex (MAPc). 10 μL MAPc samples were taken from each well and analyzed by Attune^TM^ NxT Acoustic Focusing Cytometer. The rest MAPc was resuspended in 200 μL methanol with 0.5% KOH and incubated in 37 ℃ at 250 rpm for 4 days. The mixture was then washed with methanol 2 times and diethyl ether 2 times and air-dried to give arabinogalactan-peptidoglycan (AGPG). The resulting AGPG was resuspended in 200 μL deionized water and 40 μL samples were taken from each well and analyzed by Attune^TM^ NxT Acoustic Focusing Cytometer. AGPG was digested with 0.05 N H_2_SO_4_ at 37 ℃ for 5 days with shaking at 250 rpm and washed 4 times with deionized water to give insoluble peptidoglycan (PG). PG samples were taken from each well and analyzed by Attune^TM^ NxT Acoustic Focusing Cytometer. The confocal images for the isolated sacculi were taken following the methods described below. The isolated sacculi from each well in each group was then combined and lyophilized, respectively. The lyophilized sacculi was resuspended in 0.4 mL 10 mM sodium acetate pH 5 with 25 μg/mL mutanolysin and lysozyme and digested at 37 ℃ for 20 h. The mixture was then spun down and the supernatant was filtered with 10 kDa MWCO and lyophilized. Lyophilized muropeptide was dissolved in 30 μL MilliQ water and an aliquot was applied to a C18(2) column (Luna 5 µm 100Å 250 x 4.6 mm) connected to an Agilent LC-QTOF (Agilent 1260 Infinity II Prime LC with Agilent 6545B QTOF). Muropeptide samples were eluted with a 2 to 30% linear gradient of water to acetonitrile (0.5% formic acid) at 0.4 mL/min. The muropeptides were analyzed by MS using MassHunter software.

**Confocal imaging (for whole cell labeling)** Glass microscope slides were spotted with a 1% agarose pad and 2 μL of fixed bacterial samples were deposited onto the agarose. Samples were covered with a micro cover glass and imaged using a Zeiss 880/990 multiphoton Airyscan microscopy system (63x oil-immersion lens) equipped with a 488 nm laser. Images were obtained and analyzed via Zeiss Zen software. We acknowledge the Keck Center for Cellular Imaging and for the usage of the Zeiss 880/980 multiphoton Airyscan microscopy system (PI- AP: NIH-OD025156).

**DBCO modification of polystyrene beads** 100 μL amino functionalized polystyrene beads (5% w/v, 5 mg) were spun down at 21000 g for 10 min in a 1.7 mL ebb tube and washed with 1 mL deionized water before use. The beads were then spun down and resuspended in 1 mL 20 mM sodium borate buffer pH 9 with 1 μg/mL DBCO-NHS and reacted in 37 ℃ for 2 h with shaking. The resulting beads were spun down at 21000 g for 10 min, washed once and resuspended in sodium borate buffer. 20 μL acetic anhydride was added to the suspension and reacted in 37 ℃ for 2h with shaking. The resulting product was then spun down and washed twice with 1 mL PBS and resuspend in 1 mL PBS for further use.

**Benchmarking DBCO modified polystyrene beads with Fl-Az** DBCO modified polystyrene beads were 1 to 1 diluted in PBS before adding to the assay. To a 96-well plate added 5 μL beads each well to 95 μL PBS containing different amount of **Fl-Az** to make up the concentrations indicated in the main text. The plate was incubated in 37 ℃ for 1 h and the beads were spun down at 2700 g for 10 min. The supernatant was decanted, and the beads were washed with 200 μL PBS once and then resuspended in 200 μL PBS. The samples were then analyzed by Attune^TM^ NxT Acoustic Focusing Cytometer.

**Competition with the test molecules on DBCO modified polystyrene beads** DBCO modified polystyrene beads were 1 to 1 diluted in PBS before adding to the assay. To a 96-well plate added 5 μL beads each well to 50 μM test molecules in the mini library with a final volume of 100 μL. The plate was incubated in 37 ℃ for 2 h and the beads were spun down at 2700 g for 10 min. The supernatant was decanted and 100 μL 50 μM **Fl-Az** was added to each well. The plate was then incubated in 37 ℃ for 1h and spun down at 2700 g for 10 min. The supernatant was decanted, and the beads were washed with 200 μL PBS once and then resuspended in 200 μL PBS. The samples were then analyzed by Attune^TM^ NxT Acoustic Focusing Cytometer.

***Mycobacterium tuberculosis* labeling with TetD.** Early log-phase *M. tuberculosis* (OD_600_ 0.1) was incubated with +/- DBCO-tetrapeptide 25 μM probe for 72 hours, then washed twice with PBSTrB (PBS supplemented with 0.05% Triton-X-100^2^, 0.01% BSA) and treated with for 1 h with 50 μM **FL-az** in 7H9 medium. Cells were then washed three times with PBSTrB, fixed with fresh prepared 4% PFA for 2 hours at room temperature, washed twice with PBSTrB, and resuspended in PBS for flow cytometry analysis.

***Mycobacterium tuberculosis* PAC-MAN assay.** Early log-phase *M. tuberculosis* (OD_600_ 0.1) was incubated with +/- **TetD** 25 μM probe for 36 hours, then washed twice with PBSTrB and treated with the azide-modified test molecules 50 μM in 7H9 medium for 2 h in shaking at 37°C. Cells were then centrifuged to remove the test molecules and treated with **FL-Az** 50 μM in 7H9 medium for 1 h in shaking at 37°C. Cells were then washed three times with PBSTrB, fixed with fresh prepared 4% PFA for 2 hours, washed twice with PBSTrB, and resuspended in PBS for flow cytometry analysis.
